# Multiresolution molecular dynamics simulations reveal the interplay between conformational variability and functional interactions in membrane-bound cytochrome 2B4

**DOI:** 10.1101/2024.04.18.590076

**Authors:** Sungho Bosco Han, Jonathan Teuffel, Goutam Mukherjee, Rebecca C. Wade

## Abstract

Cytochrome P450 2B4 (CYP 2B4) is one of the best characterized CYPs and serves as a key model system for understanding the mechanisms of microsomal class II CYPs, which metabolize most known drugs. The highly flexible nature of CYP 2B4 is apparent from crystal structures that show the active site with either a wide open or a closed heme binding cavity. Here, we investigated the conformational ensemble of the full-length CYP 2B4 in a phospholipid bilayer, using multiresolution molecular dynamics (MD) simulations. Coarse-grained MD simulations revealed two predominant orientations of CYP 2B4’s globular domain with respect to the bilayer. Their refinement by atomistic resolution MD showed adaptation of the enzyme’s interaction with the lipid bilayer, leading to open configurations that facilitate ligand access to the heme binding cavity. CAVER analysis of enzyme tunnels, AquaDuct analysis of water routes, and Random Acceleration Molecular Dynamics simulations of ligand dissociation support the conformation-dependent passage of molecules between the active site and the protein surroundings. Furthermore, simulation of the re-entry of the inhibitor bifonazole into the open conformation of CYP 2B4 resulted in binding at a transient hydrophobic pocket within the active site cavity that may play a role in substrate binding or allosteric regulation. Together, these results show how the open conformation of CYP 2B4 facilitates binding of substrates from and release of products to the membrane, whereas the closed conformation prolongs the residence time of substrates or inhibitors and selectively allows the passage of smaller reactants via the solvent and water channels.

**Impact:** Our findings from multiresolution molecular dynamics simulations elucidate the structural dynamics of the membrane-bound CYP 2B4 enzyme and how these affect substrate recognition and processing, as well as inhibitor binding. As CYP 2B4 is a representative of the class of cytochrome P450 enzymes that metabolize most known drugs, the insights from this study are pertinent to the prediction of drug metabolism and the design of CYP inhibitors as therapeutics.

## Introduction

Cytochrome P450 (CYP) enzymes form a superfamily of heme-containing monooxygenases that play a crucial role in the metabolism of a wide array of both endogenous and exogenous compounds^1^. Rabbit CYP 2B4, a member of the CYP 2B subfamily, has been extensively studied as a model for mammalian xenobiotic-metabolizing enzymes since its initial isolation from phenobarbital-induced rabbit liver microsomes^2,3^. Its prominence in research stems from its utility in elucidating CYP interactions with lipids and redox protein partners, as well as in providing insights into mechanistic aspects of mammalian CYP enzyme catalysis. A hallmark of CYP 2B4 is its high conformational plasticity, which enables the enzyme to accommodate a diverse array of substrates with varying sizes, shapes, and chemical properties^4,5^. The human homologue of CYP 2B4 is CYP 2B6, which shares a sequence identity of 78.7 %^6^.

Microsomal P450s, including CYP 2B4, exhibit a conserved domain architecture characterized by a globular heme-binding domain tethered to the membrane via an N-terminal anchor. The membrane tethering facilitates the enzyme’s interaction with its lipid environment and its redox protein partners, CYP reductase (CPR) and cyt b5, which is essential for electron transfer during the catalytic cycle^7^. The membrane association also influences the enzyme’s accessibility to substrates, which may come from the membrane or from the cytosol^8^.

CYP 2B4 displays the prototypical CYP topology, composed of a single polypeptide chain that arranges into 12 principal α-helices (labeled A-L), interspersed with additional helical segments (B’, F’, and G’), and four β-sheets (numbered 1-4) (see Figure 1 for a depiction of CYP 2B4 with the secondary structures that are most important for this study). These elements form a globular domain that is tethered to an N-terminal transmembrane (TM) helix by a flexible linker^4,5^. The ligand-free crystal structure of CYP 2B4 (PDB ID: 1PO5, resolution: 1.6 Å) unveiled a large hydrophobic binding cavity in the core of the enzyme^9^. The binding site is wide open to solvent and hosts a a cysteine-bound heme cofactor, the position and coordination of which are modulated by the surrounding active site residues. These residues, along with the B’ and F’-G’ regions, are implicated in forming the substrate access channel to the active site and exhibit large conformational variations amongst different crystal structures^10−12^. The binding pocket’s conformation is dynamic, with the heme proximal side being relatively rigid and the distal side, where the substrate binds, being more flexible, allowing the enzyme to adapt to various substrate geometries^13^.

**Figure 1.**
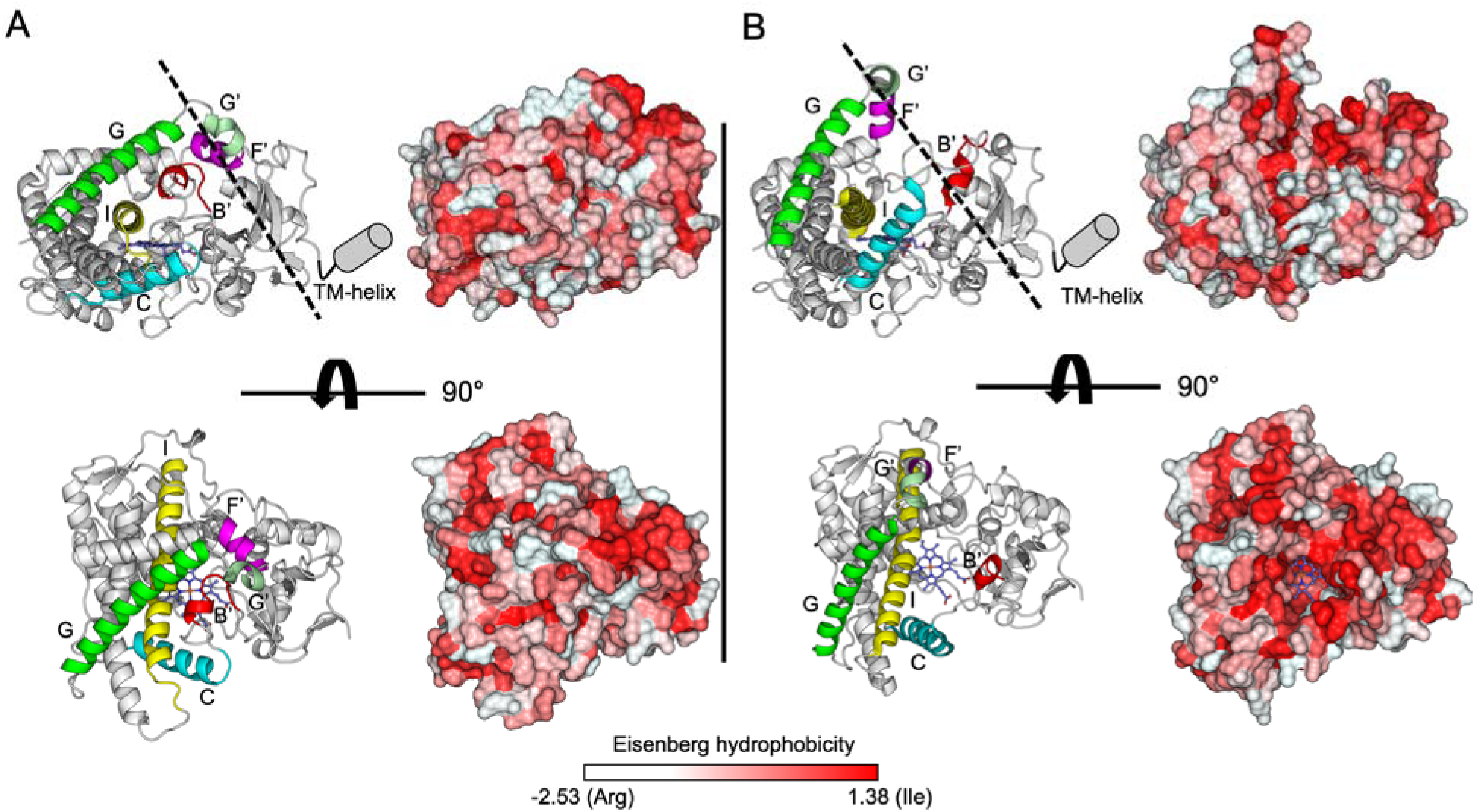
Crystal structures of CYP 2B4 in (A) closed (PDB-ID: 1SUO^15^) and (B) open (PDB-ID: 1PO5^9^) conformations. Important secondary structure elements are highlighted in color on the cartoon representations of the protein. The catalytic heme is displayed in blue stick representation. In the upper panels, the missing N-terminal transmembrane helix is indicated by a grey cylinder and the expected location of the surface of the membrane is indicated by a dashed line. These are absent from the lower panels since they show the view from the membrane. The protein surface is shown colored by hydrophobicity according to the residue-wise Eisenberg hydrophobicity scale. In the open conformation (B), a hydrophobic funnel-like opening towards the membrane is visible as a continuous red area.

The diversity of CYP 2B4’s ligand-binding capabilities is evident from available crystal structures. These structures reveal different ligand binding modes and different conformations of the globular domain, such as the binding of the antiplatelet drug ticlopidine in a hydrophobic pocket near the heme group^14^, and the closed conformation induced by the heme-coordinating inhibitor (4-(4-chlorophenyl)imidazole) (CPI)^15^. Ligand binding can also lead to conformational changes that affect the enzyme’s specificity, as seen with the calcium channel blocker amlodipine, which induces an open configuration of the B’, F’, G’, and G helices around the binding cavity^16^. The crystal structure of CYP 2B4 bound to the bulkier antifungal drug bifonazole (BIF) reveals an even more open conformation of the enzyme, with a wider active site cavity compared to other ligand-bound structures and the I-helix bent to accommodate the ligand^17^(see Fig. S9). Comparison of different crystal structures highlights the plastic regions of CYP 2B4 that undergo conformational changes upon ligand binding, showing how different substrates affect the active site cavity^17^.

The conformational plasticity of CYP 2B4 is of significant interest for xenobiotic metabolism and has implications for understanding the wider CYP superfamily, which includes human drug-metabolizing enzymes such as CYP 3A4, CYP 2D6, and CYP 2C9, which also exhibit conformational variations that contribute to their broad substrate specificity and metabolic capabilities. Understanding the conformational dynamics of these enzymes can provide insights into their individual functions and aid in the design of isoform-selective inhibitors or activators^18^. Computational approaches, including molecular dynamics (MD) simulations, have been extensively applied to study the conformational landscape of CYP 2B4 and other CYPs. These simulations have revealed the existence of multiple substrate access channels and the impact of the membrane environment on the structure and dynamics of CYP 2B4^19–22^.

The aim of this study is to provide a thorough understanding of the implications of the structural dynamics of membrane-bound CYP 2B4 for ligand access to and egress from the active site. For this purpose, we employed a multi-resolution MD simulation approach to build and simulate models of CYP 2B4 in a phospholipid bilayer in different conformational states of the enzyme and with different ligands. Initial coarse-grained (CG) MD simulations provided a broad sampling of the conformational landscape, which was then refined in all-atom (AA) MD simulations. CAVER^23,24^ and AQUADUCT^25^ analyses were carried out to characterize active site tunnels and water channels, respectively, and Random Acceleration Molecular Dynamics (RAMD) simulations were performed to identify pathways for ligand egress. Finally, simulations were performed of the re-entry of an inhibitor from the membrane into the heme binding cavity. Together, this combination of simulation and analysis techniques provides the most comprehensive understanding to date of the interplay between the conformational dynamics of membrane-bound CYP 2B4 and substrate, product, inhibitor and water passage to and from the enzyme’s active site.

## Results and Discussion

### 1. Membrane insertion CG MD simulations reveal two predominant orientations of the CYP 2B4 globular domain

CG MD simulations of CYP 2B4 were performed with the Martini 2.2 force field^26^ to efficiently sample the configurational space of the full-length protein in a phospholipid bilayer. The simulations started with either the closed or the open conformation of the globular domain resolved by X-ray crystallography (PDB ID: 1SUO, 1PO5^9,15^), see **Figure 1**. These two conformations differ most in the region distal to the heme. The F’/G’ and G-helices, along with the BC loop containing the B’ helix, are more tightly packed near the ligand binding cavity in the closed inhibitor-bound structure (PDB ID: 1SUO) than in the open ligand-free CYP 2B4 structure (PDB ID: 1PO5). The ligand-free structure has the F’/G’ helices positioned further from the I-helix and the heme, resulting in a more open heme cavity. For each simulation replica, the globular domain was initially positioned in a random orientation above the lipid bilayer and connected to the N-terminal TM helix via a 29-amino-acid residue long peptidic linker. The initial structures of the linker were assigned diverse conformations to ensure sampling of a wide range of possible arrangements of the CYP 2B4 domains and the membrane bilayer, see **Figure S1**. For each of the two structures of the globular domain, 10 replica CG MD simulations were run, each of 7 to 12 µs duration. The CG MD simulations of the closed and open CYP 2B4 structures were compared by analyzing the distribution of the two tilt angles that describe the orientation of the globular domain in the phospholipid bilayer and the axial distance between the center of mass (CoM) of the globular domain and the center of the lipid membrane, see **Figure 2**.

**Figure 2.**
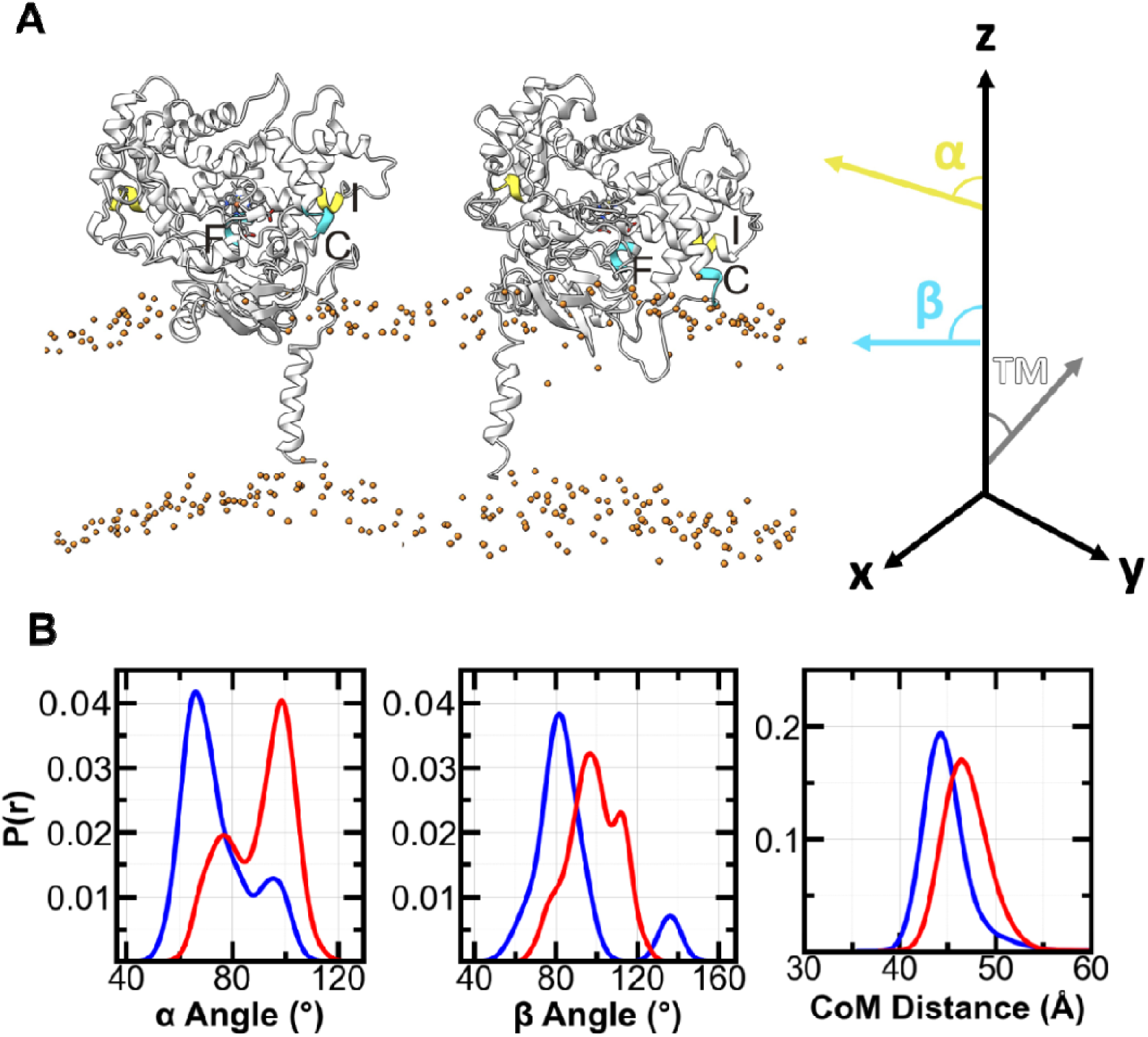
Positioning of the globular domain of CYP 2B4 with respect to the membrane bilayer in CG MD simulations. (A) Representative structures extracted from the CG simulations of closed (left) and open (right) CYP 2B4 (see Methods for details). Tilt angles are defined according to the angle between the z axis perpendicular to the membrane and pre-defined vectors: α angle, for the vector between the center of masses (CoM) of the backbone atoms of the first four and last four residues of the I-helix; β angle, for the vector between the CoMs of the backbone atoms of the first four residues of the C-helix and the last four residues of the F-helix; TM tilt angle, for the vector between the CoMs of the backbone atoms of the first and last four residues of the TM helix. Phosphate atoms of the phosphocholine are coloured in orange to represent lipid bilayer. (B) Probability densities of the α and β angles and the axial distance of the CoM of the globular domain to the center of the lipid bilayer for the converged last 2 μs of all the CG simulations for the closed (red) and open (blue) conformations of the CYP 2B4 globular domain.

For the closed conformation, the α angle, which tracks the orientation of the I-helix in the globular domain with respect to the membrane normal, converged to around 100°, with a smaller population at 75°. The sub-population of the α angle was contributed from replicas 3, 8, and 9, which also converged to relatively lower values between 80° to 100° of the β angle, measuring the orientation of the globular domain with a reference vector from the C to F helices (**Figure S2**). Overall, the β angle converged to a distribution around 100°, with a right shoulder near 115° from replicas 1, 2, and 7.

Simulations of the open structure of the globular domain revealed a tendency to lower α and β values than those for the closed structure (Figure 2). For the open conformation, the α angle converged to around 65°, with a smaller population at approximately 95°. Simultaneously, the β angle converged with a major population around 80° and a minor population at around 135°. This minor population of the β angle corresponded to the right shoulder of the α angle distribution, both originating from replica 3 (**Figure S3**). Thus, while the simulations of the open conformer converged to have nine out of the ten replicas in the distribution with α angle around 65° and β angle around 80°, the simulations of the closed conformer showed three clusters in the orientation of the globular domain on the membrane with the largest having converged α and β angles around 100° (replicas 4, 5, 6 and 10). These CG MD simulations thus indicate higher variation of the orientation of the closed globular domain on the phospholipid bilayer. One reason could be the higher axial distance of the globular domain CoM from the center of the phospholipid membrane in the CG MD simulations for the closed structure at 46.4 ± 3.4 Å, compared to 44.1 ± 3.9 Å for the open structure.

During all the CG MD simulations, the α and β angles underwent changes, showing the highly flexible nature of CYP 2B4, see **Figures S2 and S3**. The simulations for the closed conformation of the globular domain showed α and β angles converging at higher values than for the open conformation, around 100° or higher. In the closed conformation, the BC loop (containing the B’ helix) remains further from the lipid bilayer than in the open conformation, with most replicas having axial distances of the CoM of the BC loop to the lipid bilayer center that converge around 40 Å or higher. Although the distance of the B’ helix from the membrane decreases during the simulations, only the F’-G’ helices position to establish a stable direct interaction with the lipid membrane.

In one replica (replica 9, see Figure S2), however, the distance of the BC loop to the lipid bilayer decreased over time to around 35 Å, along with convergence of the α and β angles to lower values around 75°. The closed conformation can thus adopt a similar orientation of the globular domain on the membrane to the predominant orientation of the open conformation, but this is a rare occurrence. The most commonly observed membrane-bound configuration for the closed conformation is with a highly tilted globular domain with the F’-G’ helix region interacting with the membrane while the side of the protein containing the B’ helix region stays further away from the membrane surface.

For the open conformation of the globular domain, the F’-G’ helices consistently moved closer to the lipid bilayer over time, resulting in direct interactions with the lipid bilayer in most replicas and the shortest distance of the CoM of the F’-G’ helices to the bilayer center being about 25 Å. In contrast, the BC loop only interacts directly with the lipid bilayer in half of the replica simulations, but in these cases (replicas 4, 5, 7, 9 and 10 in Figure S3), the hydrophobic residues of the BC loop enter inside the upper layer of the phospholipid membrane. Insertion of the BC loop (and B’ helix) into the membrane is accompanied by the C helix orienting to be nearly perpendicular to the membrane surface and with the globular domain tilted with a lower α angle. The replicas that have a higher distance between the BC loop and the lipid bilayer (replicas 3, 6 and 8 in Figure S3) tend to have higher α and β angles of around 80° or higher. In the replica in which the BC loop is furthest from the lipid bilayer after convergence (replica 3 in Figure S3), the α and β angles were 100° and 135°, respectively. These values are more similar to those for the closed conformation and to those previously observed in studies in which the closed conformation of the globular domain of different CYPs was simulated^27,28^. However, for the set of CG simulations for the CYP 2B4 open conformation, membrane-bound conformations with lower α and β angles predominate (corresponding to a lower heme tilt angle), and these have both the BC loop and the F’-G’ helices interacting with the lipid bilayer.

Overall, the CG simulations of CYP 2B4 starting with random orientations of the globular domain at a distance from the membrane show that the globular domain consistently migrates towards the membrane surface and that this motion is accompanied by major movements of the flexible linker region. Both conformations of the CYP 2B4 globular domain show preferred membrane-interacting configurations with the F’-G’ helices in or near the bilayer surface in agreement with experimental crystallographic and H/D exchange mass spectrometry^29^ studies. These configurations could allow ligand access into the heme binding cavity from the phospholipid membrane. As the CG simulations with the open and closed conformations showed distinct membrane-bound configurations, dependent on their globular domain conformations, but also showed the ability to adopt the membrane-bound orientation observed for the other conformation, we selected three membrane-bound CYP 2B4 configurations from the CG simulations for further simulation in atomic detail (see Methods for details). We refer to these as closed, open, and alternative-open.

### 2. All-atom MD simulations of closed, open, and alternative-open states of CYP 2B4 reveal different interactions with the membrane

Due to the elastic network model applied on the globular domain and TM helix, their overall conformations were retained during the CG simulations. To sample the motions within the domains as well as in the flexible regions of the protein, AA MD simulations were performed starting from the three selected structures from the CG simulations – closed, open and alternative-open. Considering the converged α and β angle values, these were two structures representing the predominant arrangements observed in the CG simulations for the closed and open globular domain conformations and an additional structure (alternative-open) from the CG simulations with the open conformation that showed α and β angles between those of the predominant configurations of the closed and open conformations.

The structures of the globular domain of CYP 2B4 in the lipid bilayer environment were rather stable in the AA simulations, see RMSD values in **Figure 3** A,B, secondary structures in **Figure S2,** and 3D structures in **Figure 4** A. For all three conformers, the distance of the globular domain from the membrane agreed with experimental data on the height of CYP above the membrane^30^ and with other simulations of CYP-membrane systems on the distance of the globular domain CoM from the bilayer center^31,32^, which converged to a value of about 45 Å, see Figure 4 B and **Table S1**. The TM helix tilt angle fluctuated around 18 ± 10° (Figure 3 I), which matched the value of 17° reported from a solid-state NMR study of this region of CYP 2B4^32^. The most notable difference between the three conformations was in the local stability of the B’ and F’-G’ regions as the RMSD values of these parts of the open and alternative-open structures were in the 4 to 8 Å range whereas the closed structure showed stable B’ and F’-G’ regions with RMSD values around 2 Å, see Figure 3 C,D. The orientations of the globular domain in the lipid bilayer differed significantly between the three conformers as the closed CYP 2B4 shows elevated levels of α and β angles compared to the open and alternative-open structures, see Figure 3 G,H. The tilt of the globular domain relative to the membrane in the open and alternative-open structures remained lower, with the α angle around 70°, than in the closed form with α∼100°. The β angle was the lowest in the open form, converging at around 74°. The closed form showed a much higher β angle of 132°, and the alternative-open form converged to an intermediate value of 98°, see also Table S1.

**Figure 3.**
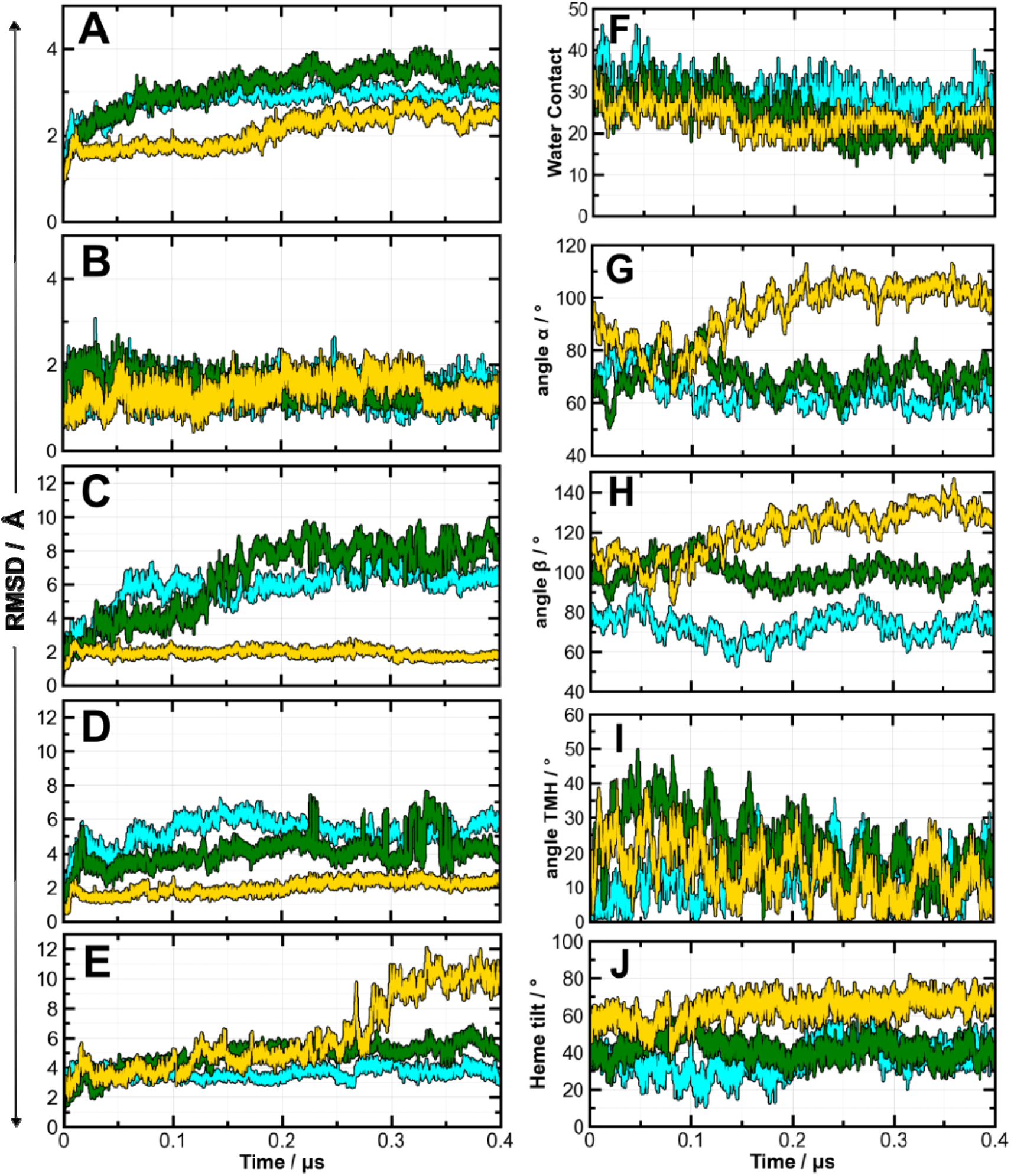
Time evolution of the structure of CYP 2B4 and the positioning of the globular domain and the TM helix with respect to the lipid bilayer during all-atom MD simulations of CYP 2B4 with three initial conformations of the globular domain: closed (yellow), open (cyan) and alternative-open (green). The Cα RMSD values, calculated with respect to the initial frame of the production run, are shown for (A) the globular domain (residues 51-492), (B) the heme+Cys436, (C) the BC loop (residues 97-118), (D) the FG loop (residues 208-229), and (E) the linker between the TM helix and the globular domain (residues 21-50). Water contacts to the heme ligand, quantified by the number of water molecules within 5 Å of the heme, are given in (F). The α (G), β (H), TM helix tilt (I), and heme tilt (J) angles are defined in Figure 2.

**Figure 4.**
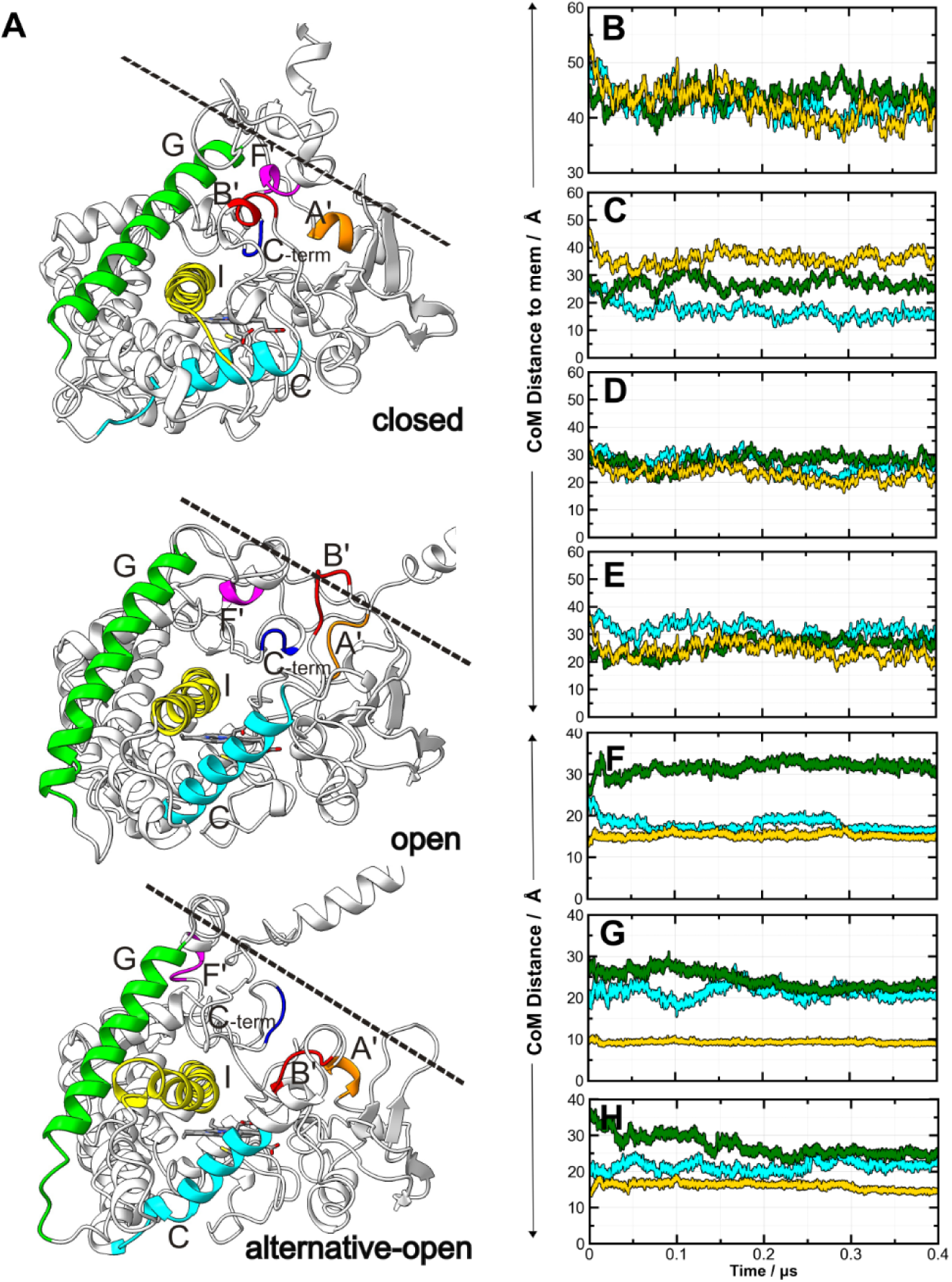
Structural adjustments of CYP 2B4 in the membrane bilayer environment during all-atom MD simulations. (A) Snapshots of the last frames of the CYP 2B4 trajectories started with closed, open, and alternative-open structures. A’ helix in orange, B’ region in red, C helix in cyan, F’ helix in magenta, G helix in green, I helix in yellow, C-terminal loop in blue. The approximate position of the membrane surface is indicated by dashed lines. (B-H) Time evolution of distances characterizing the structures during the simulations of the open, alternative-open and closed structures in cyan, green and yellow, respectively. Center-of-mass (CoM) axial distances are shown between the lipid bilayer and (B) the globular domain (residues 51-492), (C) the BC loop (residues 97-118), (D) the F’-G’ helix (residues 208-229) and (E) the linker between the TM helix and the globular domain (residues 21-50). CoM distances are also shown between (F) the A’ helix (residues 52-55) and the F’ helix (residues 213-218), (G) the B’ region (residues 102-106) and the G helix (residues 231-235) and (H) the B’ region (residues 102-106) and the C-terminal loop (residues 476-478).

Similarly to Li et al.^33^, the distances between three pairs of clusters of residues were computed to track the movements around the entrance to the active site cavity: between the A’ and F’ helices, the B’ helix region and G helix, and the B’ and C-terminal loops, see **Figure 4** F-H. The most notable difference in the alternative-open conformation was the positioning of the F’-G’ helical region with a distance between the A’ and F’ helices of around 30 Å, whereas this distance was about 15 Å in the open and closed conformations. The two monitored distance pairs for the B’ loop, to the G helix and to the C-terminal loop, fluctuated at elevated levels above 20 Å in both the open and alternative-open states, whereas in the closed state, the distances fluctuated less and were in the 10-15 Å range. Overall, the B’ loop and F’-G’ helices, located near the substrate binding pocket on the distal side of heme, had shorter distances to the respective opposing side of the pocket in the closed conformation than in the open and alternative-open conformations, which showed more flexible and diverse arrangements of these moieties.

Hydrogen-bonds between the heme propionates and the heme binding site residues in the AA MD simulations of the three conformers of CYP 2B4 were monitored with the aid of an MD-IFP analysis, see **Figure 5** and **Figure S5.** The closed conformation of CYP 2B4 displayed the most stable conserved hydrogen-bond interactions, where the heme propionate formed stable contacts with W121, R125 and H369 throughout the simulation. Other hydrogen-bonding residues include R98, to which a heme propionate established contact only after a local conformational change, and S430, which made occasional contacts. The open conformation showed the broadest range of interactions with many alterations over the course of the trajectory but maintaining stable contacts of the heme propionates to H369, and relatively close proximity with occasional contacts to R98 and S430. The interaction profile of the alternative open trajectory was the most dynamic along the trajectory with the formation of stable contacts with H369, R98 and a transition to form transient contacts with S430 and W121 over the trajectory.

**Figure 5.**
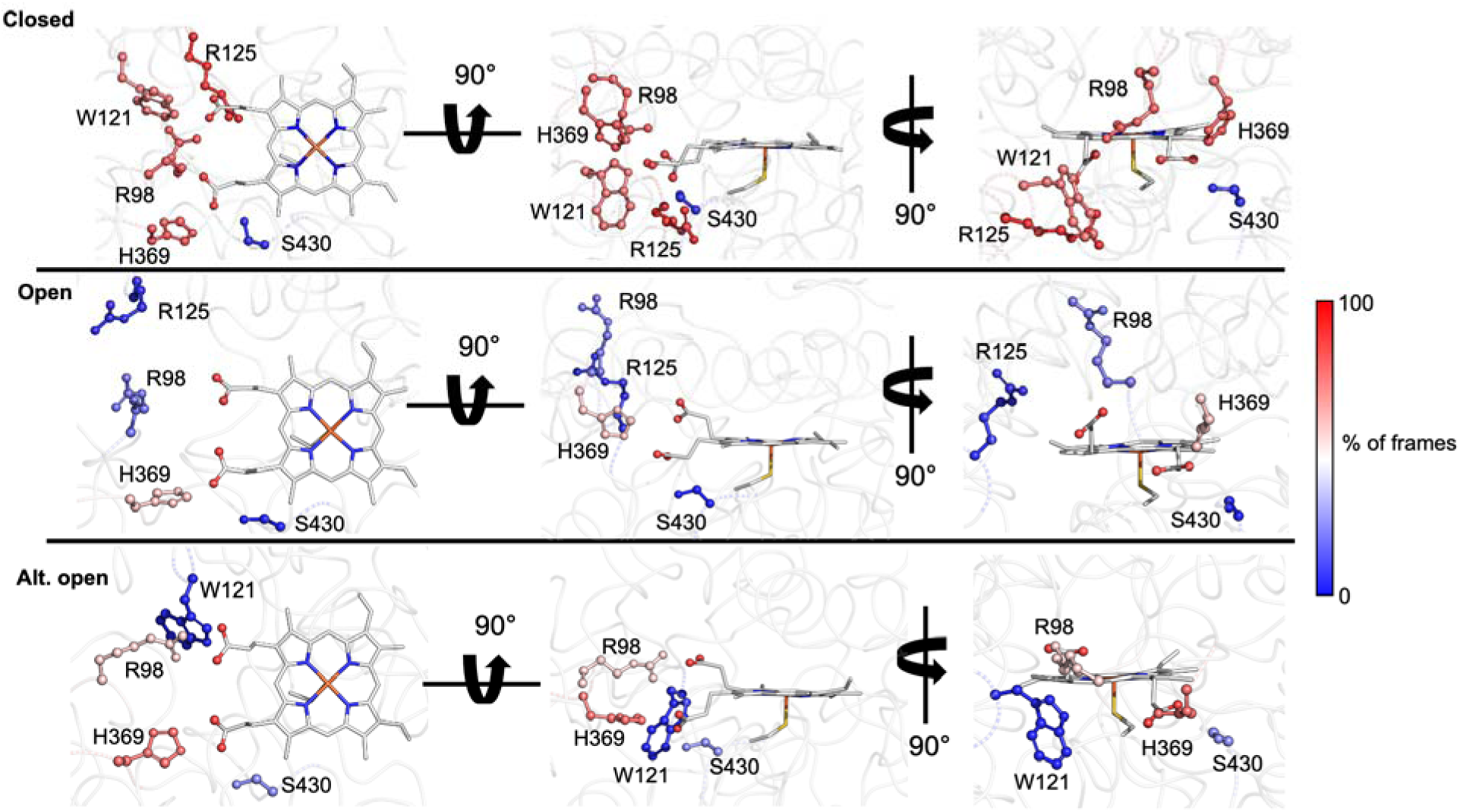
Heme-interaction patterns of the closed, open, and alternative-open states of CYP 2B4 during the AA MD simulations. Residues that form hydrogen-bond interactions with the heme propionates in more than 5 % of the analyzed frames are colored by percentage occupancy (according to a distance threshold of 3.3 Å). The heme is displayed in white carbon stick representation with the propionate oxygens as spheres.

The closed conformer showed values of the α and β angles that are within the range of those observed in our simulations of other full-length microsomal ligand-free CYPs (CYPs 2C9, 2C19, 17, 19, 1A1) in a POPC bilayer using the same CG+AA MD simulation protocol starting from crystal structures with the globular domains in a closed conformation^27,28,34^. There was also little movement in the F’-G’ region compared to the open form. However, the C-terminal end of the F helix and the N-terminal end of the F’ helix formed an interaction with the C-terminal loop that resulted in loss of helicity around the F-F’ junction. This reorientation of the F helix was accompanied by the repositioning of the G and H helices in which the H helix positioned closer to the C helix, destabilizing the C and G helices (Figure 4 A). In addition, the B’ helix and C helix stayed away from the membrane due to the B’ helix region interacting with the G and I helices. The stable interaction between the B’ loop and the G helix correlated with a noticeably lower fluctuation in the B’ loop and an RMSD of around 2 Å, compared to 7 to 8 Å in the open and alternative-open forms. The resulting globular domain configuration in the closed CYP 2B4 has a heme tilt angle of 67.9 ± 4°, which is higher than for the open and alternative-open structures (Figure 3 J). Furthermore, it is also somewhat higher than obtained in simulations of other CYPs for which the heme tilt angle ranged from 40 to 61° ^34^ and it is also higher than the values of 57-62° measured by linear dichroism for CYP 17, CYP 19 and CYP 3A4 in POPC nanodiscs^31^. A reason for the higher tilt angle could be the tight packing of the B’ helix against the G-helix above the membrane in the closed conformation of CYP 2B4.

In the open CYP 2B4 state, the B’ loop immersed into the phospholipid bilayer while bringing the adjacent C helix along, which shifted the end of the I helix neighbouring the C helix to a position closer to the membrane surface to accommodate a globular domain orientation with a low heme tilt angle of 39.9 ± 4° (Figure 4A, Figure 3J). This is at the lower end of the range observed previously in similar simulations of other CYP isoforms^34^, none of which had such an open active site as these conformations of CYP 2B4. Cojocaru et al.^28^ previously observed that CYP 2C9 tends to adopt a lower heme tilt angle when an F-G loop is present rather than the F’-G’ helices. Both the F’ and the G’ helices are present during the simulations of the closed conformation but the G’ helix unwinds during the simulation of the open conformation of CYP 2B4, which might contribute to its low heme tilt angle. A similar but less pronounced trend is visible in the alternative-open conformation. The heme tilt angle for the open conformation is also lower than the heme tilt angle obtained by Berka and co-workers for the six major human drug-metabolizing CYPs in a DOPC bilayer in AA MD simulations employing a different protocol which gave heme tilt angles ranging from 55 to 72°^35^. The crystallographic structure of ‘open’ CYP 2B4 (1PO5) is stabilized by the FG-loop region of a symmetry-related neighbor that enters into the ‘funnel-like’ opening surrounding the active-site^9^. To rule out simulation artifacts caused by removing this interacting molecule, we compared the final structure from this simulation to the crystallographic structure, which aligned well with a soluble-domain RMSD of 2.29 Å. Over the course of the simulations, the F’ and G’ helices came closer to the BC-loop region. Such a repositioning of the FG-loop is not unexpected as the ‘funnel’ in the crystal structure (1PO5) forms a large, hydrophobic surface area (see Fig. 1), indicating that closer packing of the adjacent secondary structure elements upon removal of the symmetry-related protein monomer is reasonable

In the alternative-open CYP 2B4 state, the heme tilt and α angles resembled those of the open conformation, whereas the β angle stabilized at a value between that of the open and closed conformations. While the B’ loop stabilized in a position closer to the membrane than in the closed form, the F’-G’ helices did not approach the membrane as much as in either the closed or the open conformations. Furthermore, the distance of the globular domain CoM to the membrane center was highest in the alternative-open conformer. This was because the linker between the TM-helix and the globular domain was located at the active site cavity-membrane interface, thereby separating the residues at the entrance to the active site cavity as discussed above. In addition, the alternative-open conformer showed an extensive kink in the middle of the I-helix and an unusual position of the TM helix adjacent to the globular domain. This apparent strain indicates that it would be possible for the globular domain to reorient on the membrane to allow re-positioning of the linker loop away from the active-site cavity pocket. It is also possible that the lipid-bound orientation of the protein from the CG simulation is a result of the Martini 2 force field’s tendency to overestimate protein-protein interactions^36–38^. Although it is unclear whether this unusual positioning of the loop serves any physiological relevance, this alternative-open conformation shows the transient nature of the CYP 2B4 intermediates between closed and open states.

The position of the globular domain above the lipid bilayer in all the AA MD simulations was consistent with atomic force microscopy measurements with ligand-free CYP 2B4 reconstituted in rHDL, from which the protein height was reported to be above the membrane of 35 ± 9 Å ^30^, which corresponds well to our simulation result of ∼41-44 Å for the CoM axial distance. The TM helix tilt angle for the closed, open, and alternative open conformers was 10.1° ± 4.8°, 14.1° ± 6.1° and 15.8° ± 4.9°, respectively, which is similar to the value of 17° ± 3° measured by solid-state NMR for CYP 2B4 reconstituted in DLPC/DHPC bicelles^32^.

As captured by the lower α, β and heme tilt angles in the open and alternative-open forms and the higher values in the closed forms of CYP 2B4, the degree to which the B’ and F’-G’ helices interact with lipids play a role in determining the overall tilt of the globular domain toward the membrane. More hydrophobic patches are visible in the crystal structure of the open form (Figure 1B) because the heme and active site cavity are exposed towards the membrane. Comparison of the BC loop sequence of CYP 2B4 to selected drug-metabolizing CYPs, shows that CYP 2B4 has fewer hydrophilic residues and a more clustered distribution of hydrophobic residues in this region (**Figure S6**), which could contribute to the BC loop establishing more stable lipid interactions. As a result, the orientation obtained in the open conformation places the active site cavity toward the lipid bilayer, which could allow for the entry of lipid-resident substrates.

Furthermore, it can be expected that the positioning of CYP 2B4 in membranes will be affected by substrate, favoring the closed active-site conformation, and redox protein partner binding. Indeed, in MD simulations of membrane-bound CYP 1A1 starting from a crystal structure with a closed conformation, we observed a reduction in heme tilt angle upon complexation with CPR to approximately 40° ^27^, and NMR studies and MD simulations have shown stabilizing interactions between the CYP 2B4 and cyt b5 TM helices, affecting their orientation in the membrane^39^. In addition, the CYP positioning may be affected by interactions with different components of biological membranes as CYP 2B4 has been reported to interact with sphingomyelin and induce lipid raft formation in nanodiscs^40^.

### 3. Analysis of tunnels and water paths reveals the altered specificity of functional routes for solvent molecules in different conformational states of CYP 2B4

To analyze protein tunnel formation in the simulated systems, we applied the CAVER software^23,24^. This analysis was only possible for the closed conformation of CYP 2B4 as CAVER was not able to correctly define the surface of the wide-open active site in the open structure. In the closed conformation, only the solvent channel (following the nomenclature of Cojocaru et al.^12^) was present in all the analyzed frames. The corresponding ‘solvent’ pathway was also observed as a minor pathway in the RAMD simulations for closed CYP 2B4 (see section 4). Its bottleneck radius of 1.416 ± 0.003 Å and length of 22.4 ± 0.6 Å indicate a relatively long, narrow tunnel. This shape, together with its exit towards the aqueous solvent (see **Figure 6A**), suggests it could be suited to the passage of very small molecular species, like water, ions or oxygen, rather than substrates, inhibitors or products (see **Table S2** for details).

**Figure 6.**
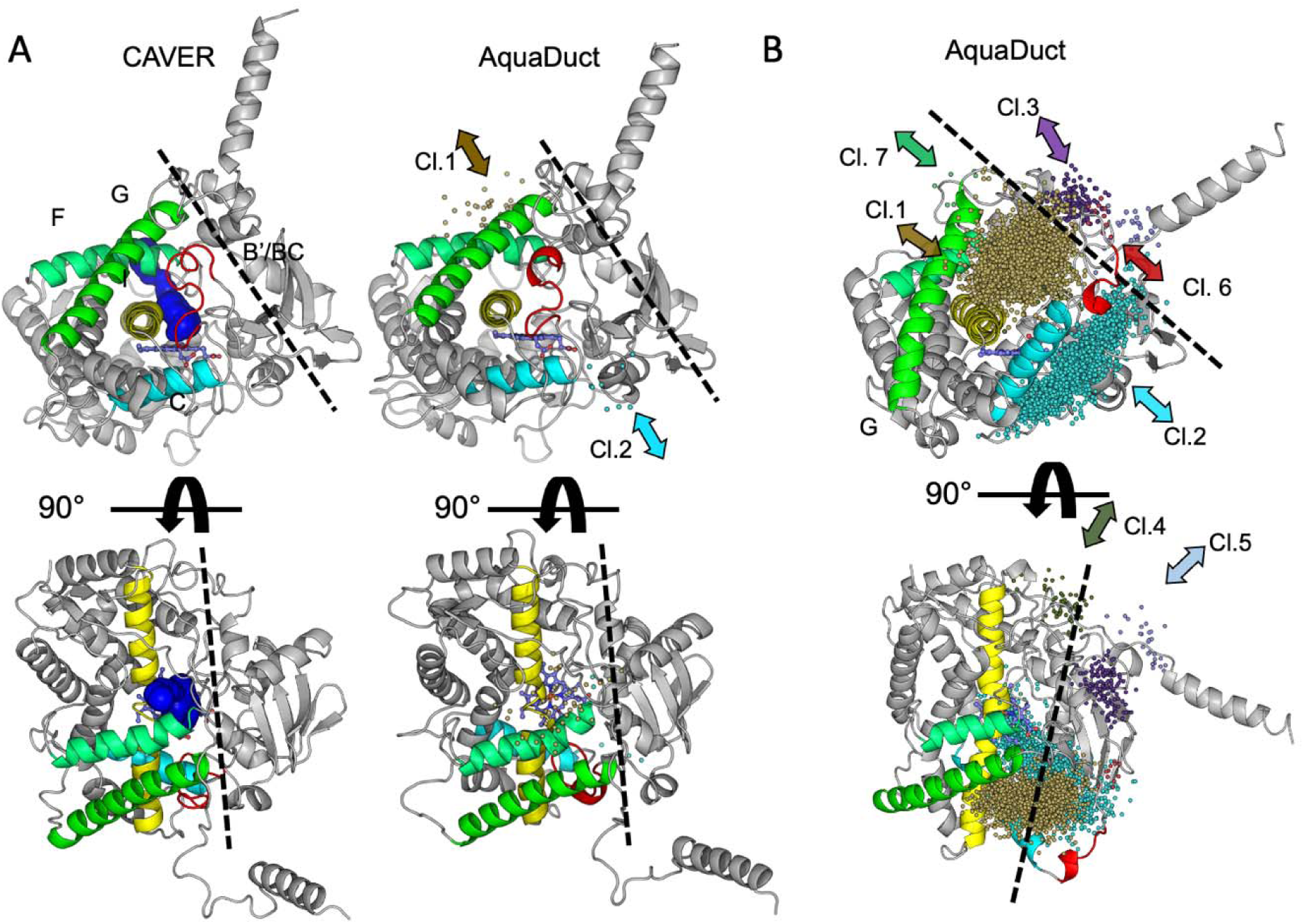
Open tunnels leading to and from the active site of CYP 2B4 in the conventional AA MD trajectories of the closed (A) and open (B) conformations as identified by analysis with CAVER and AquaDuct. The solvent channel detected by CAVER is shown by blue probe spheres in the protein structure from the final frame of the trajectory and its properties are given in Table S2. The exit points of water routes detected with AquaDuct are displayed as spheres on the protein structure from the initial frame of the analysed part of the trajectory. The spheres are colored by cluster (Cl.) number and clusters are numbered by size with Cl. 1 being the largest, see Table S3 for details. The directions of the routes followed by the clusters are indicated by arrows. The approximate position of the surface of the membrane is shown by a dashed line.

To identify routes taken by water molecules and tunnels in the simulations of the closed and open conformations of CYP 2B4, we tracked water molecules entering and exiting the convex hull of CYP 2B4 using the AquaDuct software^25^. For the trajectory of the closed CYP 2B4, results consistent with those of CAVER were obtained. The majority (Cl 1., 65%) of surface entry and exit points (referred to as ‘inlets’ by AquaDuct) of water molecules that pass through the active site belong to a cluster corresponding to the solvent channel, see Figure 6A. Thus, the hypothesis that this channel is mostly utilized by water molecules is supported by our simulations. A second cluster of openings was detected at a much lower size (Cl. 2, 14%) corresponding to the ‘water’ channel (see **Table S3** for full results).

The overall count of water molecule passages through the protein (‘separate pathways’ in AquaDuct) in the trajectories increased almost forty-fold from 56 for the closed conformation to 1997 for the open conformation. This increase indicates that the active site cavity is in constant exchange with the solvent in the open CYP 2B4. The dominant entry-point clusters in the trajectory for the open conformation correspond to pathways 2c/2ac (Cl.1, 48.6%), the water channel (Cl.2, 44.2%), channels 2f (Cl3. 3.8%) and a channel similar to pathway 5 (Cl.4, 1.5%), see Figure 6B.

These results highlight how different the closed and open conformations of CYP 2B4 are regarding solvent accessibility and exchange. Previous analysis^12^ of three crystal structures of CYP 2B4 with CAVER showed how the opening of CYP 2B4 to an intermediate structure (PDB ID: 2BDM) results in an increased count of tunnels and that there is a merging of the pathways 2a, 2ac and 2c in the open structure (PDB ID: 1PO5), which is held open by interaction with a second CYP 2B4 molecule in the crystal that coordinates the heme with a histidine. Our analysis of tunnels and water routes using CAVER and AquaDuct likewise shows that the open conformation of CYP 2B4 is highly permeable, allowing significant solvent exchange with the active site via several routes, potentially enabling the entry of small ions or molecules such as oxygen. In contrast, the observations with both CAVER and AquaDuct for the closed conformation (lower accessibility and almost exclusively via the solvent channel), indicate a more selective route for passage of small molecules. Considering that of the two simulated crystal structures, only the closed conformation contains a ligand at the required location in the active site for catalysis, the opening of a rather rigid, narrow, tunnel to the solvent could channel entrance and exit of reactants like water, protons, or oxygen, upon substrate binding, which may be important for catalytic efficiency and specificity.

### 4. Different preferred routes for ligand egress from the closed and open states of CYP2B4

We performed a total of 195 Random Acceleration Molecular Dynamics (RAMD) simulations to further investigate how the conformation of CYP 2B4, either closed or open, impacts ligand release from the active site. For this, we initially selected benzphetamine (BZP), a weight-loss causing drug and substrate of CYP 2B4 that is commonly used as a reference substrate in biochemical studies of CYP 2B4^41^, which we docked in the CYP 2B4 active site (see **Table S4** and **Figure S7**). We then selected its N-demethylated product of CYP 2B4-mediated turnover norbenzphetamine (NZP) to investigate whether the egress routes and mechanisms differ between a substrate and a product of CYP 2B4. Both compounds are relatively small and displayed a high mobility within the active site of open CYP 2B4 during bound-state simulations, with BZP almost leaving the binding site and attaching to the G helix surface. We therefore decided to also simulate the bulkier antifungal drug and monooxygenase inhibitor bifonazole (BIF), which displays specific interactions with the active site as well as imidazole-heme coordination in the crystal structure of its complex with CYP 2B4 determined by Zhao et al.^17^.

Due to the application of a randomly oriented force to the CoM of the ligand, the RAMD approach is aimed at probing the enzyme for tunnel or channel opening induced by an accelerated small molecule, rather than sampling all the dynamic motions associated with the unbinding process. 30 or 45 RAMD simulations were carried out per system. To achieve appropriate sampling of the egress routes of BZP and NZP within a maximal RAMD trajectory length of 4 ns, the random force magnitude had to be 2-fold higher for the closed than the open CYP 2B4 conformation. This indicates that the compounds have a longer residence time in the closed conformation, even though the percentage of RAMD trajectories in which no egress was observed is higher in the open than the closed CYP 2B4 conformation (see **Figure 7**A, **Table S5**). The distribution of egress routes differs markedly between the two conformational states but not between the substrate BZP and product NZP. This suggests that the ligand entry and egress route is determined largely by the conformational state of CYP 2B4 rather than the reaction stage of the ligand (substrate/product). This observation is further supported by the results for the inhibitor BIF, for which RAMD simulations were only carried out for the open CYP 2B4 conformation due to its bulkier structure than BZP or NZP. Despite requiring a higher random force magnitude for egress (8 rather than 6 kcal/mol/Å), the distribution of BIF egress routes from the open conformation is similar to that of the other ligands.

**Figure 7.**
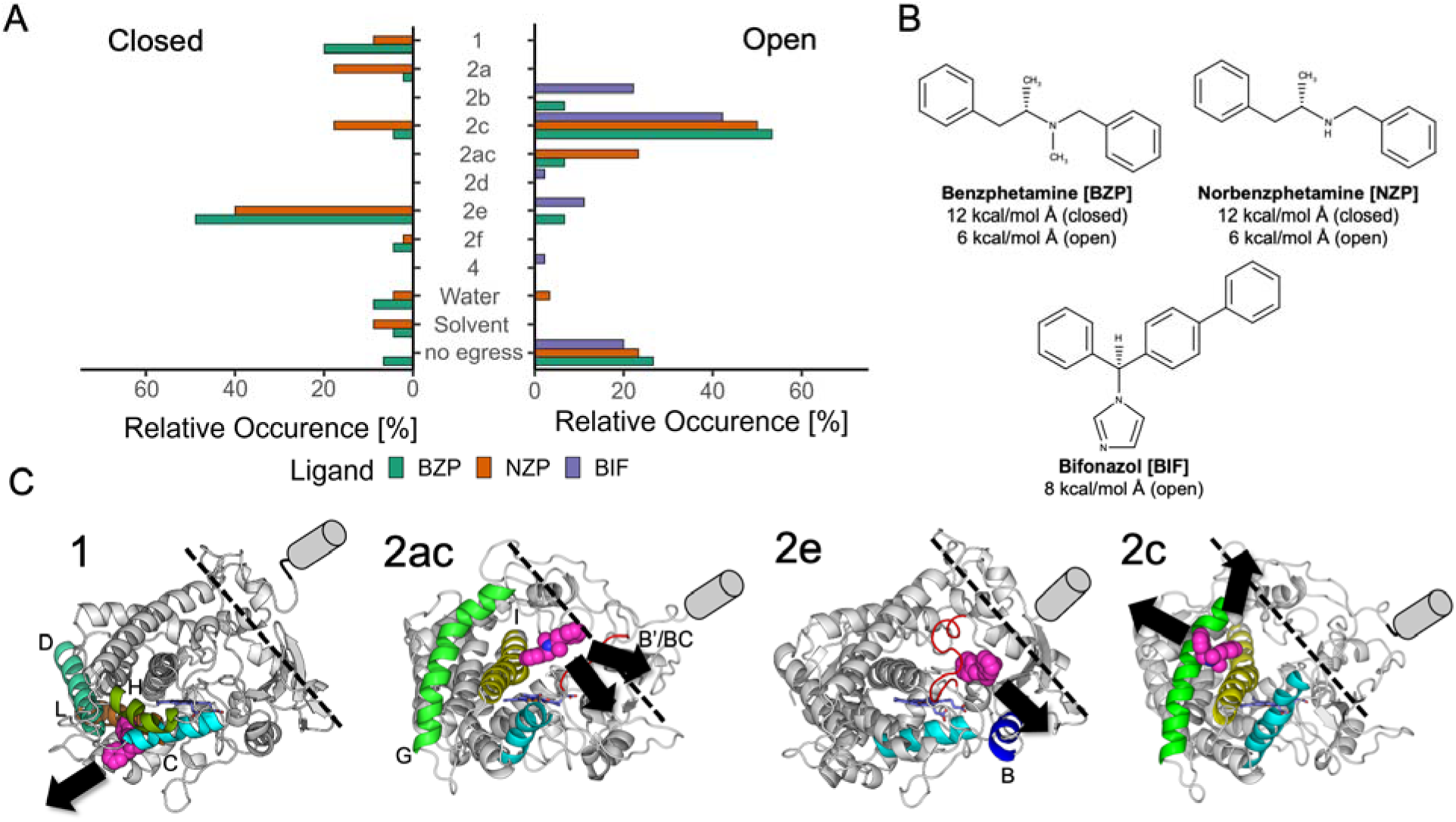
Distribution of egress routes from the active site for benzphetamine (BZP), norbenzphetamine (NZP) and bifonazole (BIF) in RAMD simulations starting from the last snapshots of conventional MD simulations of closed and open conformations of CYP 2B4. (A) Relative occurrence of each pathway over all the trajectories. (B) Chemical structures of the three ligands and random force magnitudes applied during the RAMD simulations. (C) The most common egress routes. Snapshots of egressing BZP molecules are shown as magenta van der Waals spheres with black arrows indicating the direction of egress. Pathways 1 and 2e are shown for the closed conformation of CYP 2B4 whereas pathways 2ac and 2c are shown for the open conformation. The protein is shown in cartoon representation with flanking secondary structure elements identifying the pathways colored and the TM helix is indicated by a cylinder. The approximate position of the surface of the membrane is shown by a dashed line.

For the open conformation, the dominant route for all three compounds is pathway 2c, passing through the interface between the BC loop and the I and G helices, and leading to ligand exit close to or into the phospholipid bilayer. Pathway 2ac was also often observed and was spatially partially overlapping with pathway 2c due to the rearrangement of the BC-loop region in the open conformation relative to the closed conformation. While the merging of channel 2 subclasses in ‘open’ CYPs has previously been described^42^, we distinguished between pathways 2c and 2ac in these simulations by assigning those in which the ligand formed transient contacts with the I-helix during egress to pathway 2c. The highly hydrophobic nature of the three compounds (QPlogPo/w = 3.8 for BZP and NZP, and 4.8 for BIF) indicates that they tend to be located within the microsomal membrane rather than in the cytosol. Thus, the observed egress via pathway 2c or 2ac towards the membrane indicates the likely dominant way by which substrates and inhibitors enter and products leave the active site of CYP 2B4.

For closed CYP 2B4, in contrast, BZP and NZP tended to egress mostly via pathway 2e, going through the BC loop and exiting above the membrane surface. While egress via pathway 2c towards the membrane was occasionally observed, a significant portion of the trajectories had ligands exiting via pathway 1 on the membrane-opposed face of the protein. The broader distribution of egress routes from the closed conformation, combined with the need for a higher magnitude of the random force and the lack of possibilities to enter and exit from/towards the membrane, indicate that directed and specific substrate uptake and product release to/from the membrane occur predominantly in the open conformation of CYP 2B4. Conversely, adoption of the closed conformation may serve to increase ligand residence time and retain the substrate or inhibitor in proximity to the catalytic heme.

### 5. Simulation of re-entry of bifonazole into CYP 2B4 reveals binding to an additional transient subpocket in the active site cavity

The observation of a generally high motility of the ligands within the binding site of the open conformation of CYP 2B4 during the conventional MD simulations led us to simulate the re-entry of a compound into the active site of CYP 2B4 after almost complete egress in a RAMD simulation. Considering that in open CYP 2B4 the surface lining the channels 2c/2ac leading towards the catalytic heme has large patches of hydrophobic residues (see Figure 1), we selected the inhibitor BIF for most of these re-entry simulations due to its longer residence time in RAMD simulations, its bulky nature, and its high hydrophobicity (see **Table S6)**.

As a starting point for subsequent, conventional MD simulations, a frame was selected from near the end of a RAMD simulation trajectory (at 1.288 ns before full egress at 1.374 ns) in which BIF exited via channel 2c when all but a single, unspecific contact with residue K225 had been lost and the compound had started to immerse into the phospholipid bilayer. From this starting position, three trajectories with randomly initiated velocities were generated. In two of these, BIF moved back into the funnel towards the active site. However, even after simulating each of these two replicas for 770 or 834 ns, respectively, BIF did not reach a position adjacent to the heme. Instead, in both replicas, BIF moved to a different position located between the I-helix on one side and the F- and G-helices on the other side, where it remained for at least the remaining ∼720 ns of the trajectories, see **Figure 8** A. In the third simulation, there was complete unbinding of BIF after 60 ns. In addition, in several other simulations starting with a ligand position from the end of a RAMD egress trajectory, no re-entry was observed (see Table S6). Nevertheless, the observation of two ligand re-entries supports a 2-way route for substrate access and product egress via pathway 2c, although it does not exclude the use of separate 1-way routes for substrate entry and product release^43^.

**Figure 8.**
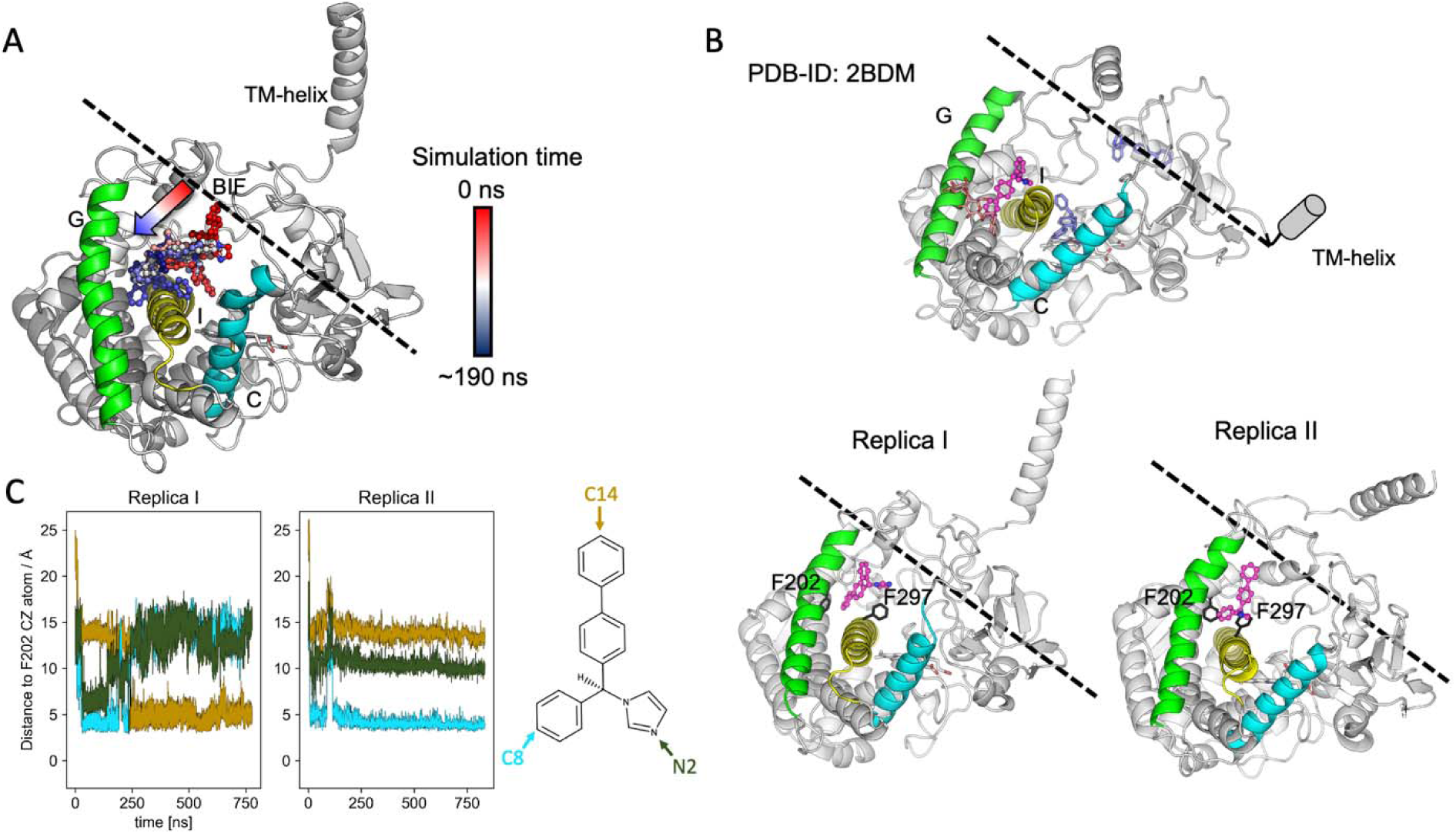
Conventional AA MD simulation of the re-entry of bifonazole into CYP 2B4 resulting in binding at an additional transient subpocket. (A) Depiction of the reentry process from superimposed simulation frames from the start of the Replica I simulation. BIF is colored according to time. Later frames, displaying the internal reorientation of BIF within the subpocket, are not shown. (B) Comparison of the positions of BIF in the last frames of the two simulations in which BIF reentered CYP 2B4 and a crystal structure of CYP 2B4 in an intermediate conformation (PDB ID: 2BDM) with three BIF molecules bound at different positions. The simulated BIF molecules and the BIF in the crystal structure in the position of interest are colored magenta. A Cymal-5 compound interacting with this BIF in the crystal structure is colored pale pink. The other two BIF molecules in the crystal structure are shown in blue. The protein is shown in cartoon representation with secondary structures of interest colored and labeled. Residues F202 and 297 are explicitly displayed in stick representation. The approximate position of the surface of the membrane is shown by a dashed line. (C) Reorientation of BIF in the additional transient subpocket during the simulations as monitored by the distances between the three ‘edge’ atoms of BIF: C8, C14 and N2, and the CZ atom of F202, a residue deep inside the subpocket. A dynamic depiction of the egress and subsequent reentry is provided by the **Supplementary movie**.

The additional subpocket observed in the BIF re-entry simulations was not present in the initial protein structure and its formation required reorientation of several residues including F296 (I-helix), which rotated towards the active site, H172 (E-helix), and L201 and F202 (both in the F-helix). Notably, the opened subpocket is more hydrophobic and tightly packed than the region next to the heme (see **Figure S8**) and can thus coordinate the hydrophobic moieties of BIF better than it. From a superposition of all 21 available crystal structures of CYP 2B4 and related proteins using the PDBe-KB website^44^, this subpocket is only occupied in one structure (PDB ID: 2BDM^17^), which has a relatively open, intermediate conformation without contact between the B’ and G helices, and in which three BIF molecules are bound to each CYP 2B4 in the crystallographic unit cell. In addition to one BIF molecule in the canonical active site, there are two alternative binding sites that were suggested to be crystallographic artefacts due to the high inhibitor concentrations used for crystallization, see Figure 8 B. One of these sites is located at the interface between two crystallographic unit cells and is at the same location as the poses identified in our simulations. Thus, the present simulations indicate that this subpocket can be formed under non-crystallographic conditions and that it could be transiently occupied upon ligand binding.

A comparison of the last frames of the two simulations showing re-entry with this crystal structure reveals that the orientation of BIF varies. It inserts in the interface between the F, G and I helices with its biphenyl moiety in the crystal structure and also, after some reorientations, in the Replica I simulation. On the other hand, it inserts stably with its monophenyl moiety in the Replica II simulation (see Figure 8 B). During the initial binding of BIF to the subpocket in Replica I, a similar pose to that in Replica II was generated yet the inhibitor remained very unstable and, eventually, reoriented to bury the biphenyl instead (see Figure 8 C). The apparent difficulty of opening the transient subpocket wide enough for the biphenyl moiety in Replica II is indicative of tighter packing of the two helices in the simulations in the absence of the Cymal-5 detergent crystallization aide, of which one molecule is located near the G helix and forms additional contacts with BIF in the crystal structure. The conformation of the subpocket between the final frame of Replica I and the crystal structure also differs and the pocket is more open in the crystal structure with BIF located deeper inside. Thus, the orientation of BIF at this site in the crystal structure may be influenced by the Cymal-5 compound and by the formation of contacts to the protein in the neighbouring unit cell. However, it is also possible that the interface region between the helices may widen on timescales beyond those of our atomistic simulations.

Gay et al. compared the crystal structures of the complexes of CYP 2B4 with three differently sized inhibitors, including BIF, identifying major structural rearrangements of the protein, including in the region of the additional subpocket^45^. Remarkably, our simulations show the formation of this subpocket by an induced-fit mechanism during simulations that were based on the open conformation of ligand-free CYP 2B4 (PDB-ID: 1PO5) in which the subpocket region between the F, G and I-helices is completely closed. Unlike the crystal structure of the BIF-CYP 2B4 complex, the simulated protein did not display a rotation of the I-helix upon BIF binding but was still able to accommodate BIF in the additional subpocket indicating that this pocket is not only present in the BIF-bound intermediate conformation in the 2BDM crystal structure.

The agreement between the crystal structure of an intermediate state of CYP 2B4 (PDB: 2BDM), which has BIF bound at this subpocket, and the simulations suggests that this subpocket may act as a transient binding site or participate in allosteric regulation. Binding at this region could be part of the substrate uptake process, with the substrate waiting in this position until the closing motion of the protein brings it towards the heme for reaction. Alternatively, it is possible that binding of a compound at this site has an allosteric role, reducing the solvent-filled volume of the active-site access tunnel and providing additional hydrophobic contact points for a second molecule entering the active site from the membrane. To test the latter hypothesis, we performed further conventional MD simulations with one BIF molecule in the additional pocket and a second BIF molecule positioned as for the re-entry simulations described above at the opening of channel 2c (data not shown). In almost all cases, the second BIF remained at the entry-site forming hydrophobic contacts with the BIF inside the additional subpocket, indicating a stabilizing effect of the inhibitor in the additional subpocket on the second entering inhibitor molecule.

Cooperative mechanisms of CYP catalysis have been widely discussed, most notably for CYP 3A4^46^. Simultaneous binding of multiple ligands, not only to the active site but also to additional sites, may assist in keeping the substrate in position for turnover by enhancing packing in the wider and less specific binding sites of more promiscuous CYP isoforms^46^. An additional ‘peripheral’ pocket in CYP 2B4 adjacent to the subpocket identified here, also located behind the F and G helices, has been investigated by Jang et al.^47^, who found that mutation to tryptophan of F202 or I241, which form the inner surface of this pocket and the additional subpocket at which BIF binds in our simulations, led to impaired turnover of 7-ethoxy-4-(trifluoromethyl)coumarin and 7-benzyloxyresorufin. A third mutation, F195W at the entry of the ‘peripheral’ pocket but far away from BIF in our simulations, had no such effect. Thus, a role of the additional subpocket occupied by BIF in our simulations in CYP 2B4 catalysis seems likely. However, Zhao et al. did not detect binding of multiple molecules of BIF to CYP 2B4 by isothermal titration calorimetry, leading them to surmise that the two additional sites in the crystal structure (PDB-ID 2BDM) that are not located in the active site are low-affinity binding sites for BIF.

Thus, it is possible that the observed binding of BIF to the additional subpocket in CYP 2B4 may not represent a mechanism in which multiple ligands bind simultaneously but rather the first step in a complex ligand-uptake mechanism. For instance, Isin et al. proposed a three-step substrate binding process for CYP 3A4 in which an initial noncatalytic ‘encounter’ complex between the substrate and enzyme is rapidly formed and this is then followed by conformational transitions relocating the substrate closer to the heme and facilitating catalysis^48^. Similar results have been obtained by Davydov and colleagues for bacterial P450eryF with the fluorescent substrate Fluorol-7GA using FRET and pressure perturbation^49^. The observed rapid association of BIF from a membrane-bound exit/entry point to the additional subpocket followed by relatively stable occupancy of this binding mode over 700 ns of simulation time points towards a similar mechanism in CYP 2B4.

More recently, Hackett simulated entry of the substrate testosterone from a membrane into membrane-bound CYP 3A4 using accelerated MD and adaptive biasing force simulations. Ligand ingress via pathway 2 (2a, 2ac or 2d/f) was observed with a 2-step entry process with an intermediate state where testosterone was stabilized by aromatic residues along the entry pathway located between the I-helix and the FG and BC loops^50^. Furthermore, it was suggested that both the intermediate and active site poses may be simultaneously occupied upon high substrate concentrations. This study also identified F304 in CYP 3A4 to be important in formation of the metastable state. This residue has previously been found to be important for cooperativity in CYP 3A4^51^ and is homologous to F297 in CYP 2B4. A conformational change of this residue is essential for the observed opening of the additional subpocket in our simulations. Amongst the major drug-metabolising CYPs (1A2, 2A6, 2C9, 2D6, 2E1 and 3A4), this position is only conserved in CYPs 2C9 and 3A4, but it is also present in CYP 2B6, the human homologue of CYP 2B4. While the study by Hackett was done on a different CYP isoform in a different (closed) conformational state and depicted a different entry channel, the concept of a two-step entry mechanism starting with a membrane-bound ligand is similar.

F297 in CYP 2B4 has been found to display a degree of flexibility within crystallographic structures of closed CYP 2B4 depending on the ligand bound in the active site^13^. The F297A mutation resulted in roughly 1.6 and 1.2-fold decreases, respectively, of the K_m_ and k_cat_ values, yet no notable change in catalytic efficiency^52^. This was, however, linked to the role of this residue in coordinating the substrate in the active site in the closed conformation as a change between an inward and outward facing orientation of F297 in CYP 2B4 occurred upon opening of the peripheral pocket.

To the best of our knowledge, our simulations provide the first observation of unbiased ligand reentry into the active site from RAMD-derived egress points. In previous work applying a similar approach to investigate ligand orientations within the membrane after egress from CYP51, re-entry events were not observed^53^. This approach of simulating ligand entry in conventional MD simulations by using input structures generated from RAMD simulations can be expected to be applicable to other systems.

The entire process, depicting the RAMD-trajectory chosen as a starting point for this approach, the selected point of last contact and the subsequent reentry simulation (Replica I) are depicted in the **Supplementary movie**.

## Conclusions

Our integrative simulation study, employing coarse-grained and all-atom conventional molecular dynamics simulations complemented by Random Acceleration Molecular Dynamics (RAMD) simulations, provides a detailed picture of the conformational dynamics and ligand interaction mechanisms of CYP 2B4 immersed in a phospholipid bilayer. The results elucidate the protein conformation-dependent modulation of ligand and solvent pathways and the enzyme’s interaction with the lipid membrane, all of which are important for the enzyme’s biological function.

### Dependence on globular domain conformation of the positioning of CYP 2B4 in the membrane

The CG simulations revealed that, depending on whether it has a conformation with an open or a closed active site cavity, the globular domain of CYP 2B4 adopts one of two major orientations when interacting with the lipid bilayer, with the BC loop and F’-G’ helices playing a central role in this interaction. Furthermore, we observed minor orientations, representing intermediate orientations and the respective opposite orientation, i.e. the closed conformation of the globular domain adopting the major orientation of the open conformation and *vice versa*. Thus, in addition to one orientation for the open and one for the closed state, we selected one intermediate orientation, labelled alternative-open, from the CG simulations for the open CYP 2B4 for subsequent AA MD.

In the AA MD of the closed, open, and alternative-open CYP 2B4, the geometric parameters characterizing the position of the globular domain on the membrane span the range of membrane orientations of microsomal CYPs observed previously, likely due to the high degree of flexibility of the CYP 2B4 globular domain in which the active site conformations range from tightly closed to wide open.

### Dependence of ligand and solvent access and egress on globular domain conformation of membrane-bound CYP 2B4

We carried out a detailed analysis of protein permeability and ligand passage from the active site of the membrane-bound full-length CYP 2B4 by analysing: 1) tunnel occurrence in conventional MD simulations by CAVER, 2) passage of water molecules through preformed and induced tunnels in conventional MD simulations by AQUADUCT, and 3) ligand egress through existent and induced tunnels by RAMD simulations that enhance ligand motion and thereby accelerate ligand egress. The results show the influence of the conformation of the globular domain on the egress or both water molecules and substrate, product, and inhibitor molecules from the active site. The open state of the enzyme supports a range of ligand egress routes, including those leading directly to the lipid bilayer, indicating that the open state supports both substrate entry and product egress, as well as being highly permeable to water molecules. In contrast, the closed state prolongs the residence time of the ligands, thereby enhancing the efficiency of catalysis by maintaining substrates in proximity to the catalytic heme center and channeling the access and egress of water via the solvent tunnel.

We observed the re-entry of the hydrophobic inhibitor BIF from an exit point to an additional transient hydrophobic subpocket in the CYP 2B4 active site cavity in conventional MD simulations starting from a RAMD simulation snapshot with the ligand outside the protein. These results support ligand uptake/release by a single 2-way route and suggest a functional role of the transient subpocket, in a two-step entry mechanism or as an allosteric site. These insights advance our understanding of CYP 2B4’s functionality and have broader implications as regards the pharmacological role of CYP enzymes in drug metabolism.

## Methods

### Preparation of full length CYP 2B4 models

Full length models of CYP 2B4 were constructed using the following high-resolution crystal structures deposited in the Protein Data Bank (PDB): “open” ligand-free CYP 2B4 (PDB ID 1PO5^9^; 1.60 Å resolution; N-terminal residues 3–21 truncated) and “closed” CYP 2B4 bound to 4-(4-chlorophenyl)imidazole (PDB ID 1SUO^15^; 1.90 Å resolution, N-terminal residues 3–21 truncated). The globular domains from these experimental structures were used for homology modelling based on the Uniprot sequence P00178, in which the missing residues (1–48) were built using Modeller v9.23^54^, while the secondary structure prediction tool TMPred^55^ was used to model the missing transmembrane helix (residues 1–20). For both “open” and “closed” CYP 2B4 models, 10 different configurations were generated with varying orientation and distance to the lipid bilayer (Supplementary Figure S1). After removing the heme cofactor, each model was converted to the CG MARTINI representation after being inserted into a pre-equilibrated CG 1-palmitoyl-2-oleoyl-sn-glycero-3-phosphocholine (POPC) bilayer as described previously^27,28^.

### CG MD simulation

A total of 20 individual CG MD simulations were performed for 7 to 12 µs each, using the GROMACS^56^ v5.0.6 software package with the Martini2.2^57^ CG force field for the protein, lipid, and water, under periodic boundary conditions. Harmonic restraints were applied on the backbone atoms of the protein except the flexible linker region (residues 21–50) with an elastic force constant of 500 kJ/mol/nm^2^ and a distance cutoff of 5–9 Å to retain the overall secondary and tertiary structure during the simulations based on the secondary structure information obtained from the DSSP^58^ server. The steepest-descent algorithm was used for energy minimization for 10000 steps with a maximum force of 5 kJ/mol/nm. A short equilibration of 20 ns with a time step of 20 fs was performed at a constant pressure of 1 bar at 310 K. Temperatures of the protein with POPC and solvent with ions were controlled separately with a velocity rescale thermostat with a coupling constant of 1 ps. A Berendsen barostat was used to achieve constant pressure using alternative isotropic pressure coupling with a compressibility of 3.0 × 10^−4^ bar^-1^, a coupling constant of 3.0 ps and a reference pressure of 1 bar. The long-range non-bonded interactions were computed using a reaction field (RF) with a dielectric constant of 15. The neighbor-list was updated every 10 steps. The Verlet pairlist algorithm was used for calculating non-bonded forces with a cutoff 1.1 nm and a Verlet-buffer-tolerance (verlet-buffer-drift) value of 0.005 kJ/mol/ps. During the production simulations of around 7 to 12 μs, the pressure coupling method was switched to the Parrinello-Rahman barostat with a coupling constant of 15 ps and the non-bonded interactions were calculated using a RF with a cutoff distance of 1.2 nm.

The CG MD simulation trajectories were analyzed for convergence by computing structural angle and distance parameters to track the orientation of the CYP 2B4 globular domain with respect to the membrane during the simulations as previously described ^27,28^. Distributions were computed of the α and β angles and the globular domain CoM - membrane CoM axial distance over the last 2 μs of the trajectories, when the positioning of the CYP 2B4 in bilayer was converged. Three representative structures were selected for conversion to atomic detail for AA MD. For the closed and open conformers, the representative structures were taken from the frames in the last 2 μs of the simulations that matched the average α and β angles over this time scale from all replicas of the respective CG simulation sets, see Table S1. The alternative-open conformer representative structure was selected from the open CYP 2B4 set to have intermediate values of the α and β angles between those of the open and closed conformations. The closed representative conformer is a frame from replica 10 from the CG simulation set initiated from the closed CYP 2B4 globular domain crystal structure, frames from replica 9 and replica 8 from the CG simulation set initiated from the open CYP 2B4 globular domain crystal structure were chosen for the open and alternative-open conformers, respectively.

### AA MD simulation

The protein and lipid membrane in the three CYP 2B4 structures from the CG simulation were converted to atomic detail as previously described ^27,28^. The crystal structures of the open and closed globular domain conformations were superimposed on the converted structures to introduce the heme in the binding site. AA MD simulations were conducted using the AMBER ff14SB^59^ force field for protein and the LIPID14^60^ force field for the POPC membrane, and GAFF-parameters for ferric, low spin, hexa- (water-) coordinated heme (‘resting state’) from Harris and colleagues^61^.

The systems were immersed in a periodic box of TIP3P water molecules with 150 mM ionic concentration. They were energy minimized using AMBER20^62^ and applying harmonic restraints with a force constant decreasing from 1000 to 0 kcal/mol/Å^2^ on the heavy atoms of the protein, as described in previous studies ^27,28^. The NAMD 2.14^63^ package was used for equilibration and production simulation runs. The NPAT ensemble was used during equilibration with constant surface area, constant pressure of 1 bar, and constant temperature of 310 K. A time step of 1 fs was used for the initial 2.8 ns equilibration run with harmonic restraints with a force constant decreasing from 100 to 0 kcal/mol/Å^2^. Pressure was controlled by using the Nosé-Hoover Langevin piston method with an oscillation time of 100 fs and a damping time of 50 fs. Temperature was controlled by Langevin dynamics with a damping coefficient of 5.0 ps^−1^. The system was equilibrated further for 12.5 ns without harmonic restraints using a time step of 1 fs for the first 2.5 ns and a time step of 2 fs for the remaining 10 ns. The Nosé-Hoover Langevin piston method was used with an oscillation time of 200 fs and a damping time of 500 fs and the temperature was kept constant by Langevin dynamics with a damping coefficient of 1.0 ps^−1^. For the production simulations, the NPT isobaric-isothermal ensemble was used with a time step of 2 fs. The electrostatic interactions were calculated using the Particle Mesh Ewald (PME) method and all bonds to hydrogen atoms were constrained using the SHAKE algorithm. Temperature was controlled by Langevin dynamics with a damping coefficient of 0.5 ps^−1^ at 310 K on non-hydrogen atoms. Constant pressure was achieved by using the Nosé-Hoover Langevin piston method with an oscillation time of 1000 fs and a damping time of 1000 fs. The production runs of the three sets of simulation were analyzed using the same structural parameter values used for the CG MD trajectory analysis. The heme tilt angle was also monitored during the simulations with the tilt being the angle between the four porphyrin nitrogens defining the heme plane and the membrane normal (z axis).

### Secondary structure analysis

Residue-wise secondary structure classification was performed using the DSSP algorithm^58^ with the ‘timeline’ feature in VMD^64^. Since the assignment of the heme and covalently bound proximal cysteine to one “residue” caused this method to fail and the cysteine is located within a long loop-region, it was removed from the analysed trajectory with CPPTRAJ^65^. The analysis was then performed at time intervals of 200 ps.

### Analysis of heme interactions

Residue-wise interaction patterns of the heme and proximal cysteine were identified with the MD-IFP workflow^66^. Potential interacting residues were selected as the globular domain of CYP 2B4 and the IFP generation was set to include nonspecific interactions. The results from this analysis were combined with visual analyses of the trajectories using VMD to identify the hydrogen-bonding partners of the heme propionates. All hydrogen-bonding contacts, but not those formed by the proximal cysteine residue, were recorded and the minimal distance between any of the propionate oxygens and hydrogen-bonding competent residues of the recorded residues were computed in a separate step using the MDAnalysis library^67^. All interactions were analyzed in steps of 200 ps.

### AquaDuct and CAVER analysis

Protein tunnels in the trajectories of the closed conformation of CYP 2B4 were identified with CAVER 3.0^23,24^. Analysing the open trajectory with CAVER did not yield meaningful results as the protein surface was detected to be formed by the heme residue itself and the open ‘funnel’ leading towards the active site was classified as the protein surroundings.

For the closed conformation, a trajectory file of the last 100 ns of each trajectory with frames saved at intervals of 20 ps was prepared. Each frame of this trajectory was then aligned to the initial frame by superposition of the globular domain residues and exported as a pdb-format file for CAVER. Then CAVER was run with default parameters using a probe radius of 1.4 Å and shell radius and shell-depth of 4 Å. The starting points were defined as the heme-iron and heme-alpha nitrogen (NA) atoms.

Water routes in the trajectories of closed and open CYP 2B4 were analyzed with the AquaDuct software (v.1.0.11)^25^. Trajectory files of the last 100 ns of each simulation were generated, including water molecules, ions, and the phospholipid bilayer, with snapshots at intervals of 6 ps. These trajectories were then centered on the catalytic heme-cysteine residue with GROMACS 2020. The AquaDuct analysis was mostly run with default parameters: the ‘scope’ was defined as the globular domain of CYP 2B4 (residues 20 to 492) and the ‘object’ was defined as water molecules within 5 Å of the iron-atom of the heme. The option ‘scope_everyframe’ was inactivated due to centering issues and cluster were manually combined in a second run of the program to yield intuitive and biologically meaningful insights. An additional recursive clustering using the ‘birch’ method was utilized in the analysis of the open conformation to subdivide a large cluster into two. For this, the ‘recursive_threshold’ parameter was set to <0.5.

### Generation of ligand-bound CYP2B4 complexes

BZP was inserted into the active site of closed (PDB:1SUO) and open (PDB:1PO5) crystal structures of CYP 2B4 with Autodock Vina 1.5.6^68^. For this purpose, a 50×50×50 (closed) / 92×64×68 (open) point receptor grid with a spacing of 0.375 Å centered 7.6 (closed) / 10.6 Å (open) above the heme iron in the center of the active site was generated with the following residues set to be flexible during the docking run:

**Closed**: S96, R98, K100, I101, V103, V104, I107, F108, Q109, Y111, V113, I114, F115, F297, V367

**Open**: F297, E301, T302 I363

Autodock Vina’s ‘exhaustiveness’ parameter was set to 10, balancing computing time versus accuracy, and the maximum number of models was 5 (closed) and 10 (open).

The pose to be used in subsequent AA MD simulations was selected by requiring 1) a good docking score, 2) the reactive methyl group of BZP to be close to the heme iron, and 3) the pose to be close to the I-helix where most ligands in crystal structures of CYP 2B4-ligand complexes form contacts. Accordingly, poses #2 (closed) and #8 (open) were selected for further simulation (see Table S4 and Figure S7).

Due to the very high similarity between the two compounds, NZP was inserted into the active sites of the closed and open crystal structures by superposition on the selected binding poses of BZP. Since docking of BIF with Autodock Vina did not yield satisfactory results and due to the availability of a crystal structure of BIF bound to an intermediate conformation of CYP 2B4 (PDB:2BDM^17^), this was used as a template for the binding pose of BIF. The ligand was inserted into the open crystal structure (PDB:1PO5) by superimposing the two crystal structures by aligning the heme cofactors^19,66^.

For subsequent MD simulations, GAFF force-field parameters for the three compounds were derived with the Antechamber program in AMBER Tools^62^. RESP partial atomic charges were derived from quantum-chemical calculations using Gaussian09^69^. For this, the structures of the compounds were geometry-optimized at the B3LYP level using a 6-31G* basis set and then the Hessian was computed (by computation of vibrational frequencies using the *freq-*keyword) to rule out the identification of saddle-point geometries. Subsequently, molecular electrostatic potentials were computed for the optimized geometries at the Hartree-Fock level with a 6-31G* basis set using the *iop(6/50=1)* and *pop=mk* keywords. The docked complexes were energy minimized and then converged in short MD simulations of 100-200 ns in NAMD using the protocol described above.

### Random Acceleration Molecular Dynamics simulations of ligand egress and re-entry

The egress of BZP, NZP and BIF from the CYP active site was simulated by the Random Acceleration Molecular Dynamics (RAMD)-method using a modified GROMACS 2020 version: GROMACS-RAMD 2.0 available at: (https://github.com/HITS-MCM/gromacs-ramd). Input structures were generated by extracting the final frames from the previous MD simulations run with NAMD (see above) after 113.3 ns (BZP, closed), 116.1 ns (NZP, closed), 217.7 ns (NZP, open), and 127.6 ns (open, BIF). Since BZP displayed a very high mobility in the active site in the open conformation, an earlier frame at 20 ns, which more closely represented its binding pose, was utilized. GROMACS-compatible structures and topology files were generated by conversion using ParmEd^70^. All systems were equilibrated and subsequently sampled in two consecutive, unrestrained simulations in the NPT ensemble with a timestep of 2 fs. The Particle Mesh Ewald method using a Verlet-cutoff scheme (using a cut-off distance of 1.1 nm for the short-range neighbor list) was applied for computing long-range electrostatic interactions. Bonds to hydrogen atoms were constrained using the LINCS algorithm^71^. The first simulation (equilibration 1) utilized the Berendsen thermostat (τ = 1.0 ps) and a semi-isotropic Berendsen barostat (τ = 5.0 ps, compressibility = 4.5×10^-5^ bar^-1^) to heat the system to 310 K at a constant pressure of 1 bar. Three separate molecular groups consisting of (1) the protein and ligands, (2) the phosphatidylcholine lipid bilayer, and (3) the water and ions, were used for thermal coupling. To achieve increased sampling of the bound state, the second equilibration of 20 ns was run with randomly initiated velocities. This simulation was repeated once (open: BZP/NZP) or twice (open: BIF; closed 2B4: BZP/NZP) with modified random number generation seeds. In these simulations, a semi-isotropic Parrinello-Rahman barostat (τ = 5.0 ps, compressibility = 4.5×10^-5^ bar^-1^) and a Nosé-Hoover thermostat (τ = 1.0 ps) were utilized to equilibrate the systems to 310 K and 1 bar. Each resulting system was then subjected to 15 RAMD simulations in the NPT ensemble using identical parameters to those used in the previous simulations. The RAMD simulations were stopped after the ligand CoM had reached a distance of 60 Å from the heme-iron atom of CYP or after 4 ns of simulation time. The random force magnitude was varied according to the system simulated as follows: closed 2B4: 501 kJ/mol/nm (12 kcal/mol/Å), open 2B4: NZP: 250 kJ/mol/nm (6 kcal/mol/Å), BZP: 251 kJ/mol/Å (6 kcal/mol/Å), BIF: 334 kJ/mol/Å (8 kcal/mol/Å). The randomly chosen direction of the force was evaluated every 50 simulation steps and the direction changed randomly if the ligand did not move further than the distance-travelled criterion of 0.0025 nm. The number of RAMD simulations per system depended on the results of the first two sets of 15 simulations, each of which were started from two preceding independent, conventional MD simulations of the bound state. If more than three unique egress routes, i.e. routes which were only sampled in one of the two sets were observed, a third set of 15 trajectories was simulated. In addition, three sets of 15 trajectories were generated for BIF, despite high agreement between the first two sets, so as to yield more and diverse poses at the point of last contact for subsequent simulations of the reentry of BIF into the protein. The ligand egress routes were analyzed by visual inspection using VMD 1.9 ^64^.

Entry of BIF to CYP 2B4 was simulated by extraction of frames from the corresponding RAMD trajectories at a point where some of the last contacts were formed between the protein and ligand. These frames were then subjected to conventional MD simulations using identical parameters to those for the RAMD simulations (see above).

### Octanol-water partition coefficient predictions

QPlogPo/w values were calculated using QikProp 7.4^72^ in the Maestro environment (Schrödinger release: 2022-r4). Prior to the calculation, potential protonation states at pH 7.0 ± 2.0 were generated with LigPrep. Subsequently, QikProp was run with default parameters. If multiple protonation states were predicted, the QPlogPo/w values were averaged.

## Supplementary Material

PDF file containing Tables S1-S5 and Figures S1-S9, which provide further details of the results.

Movie depicting egress of BIF in a RAMD simulation, the selection of a starting point from this simulation and subsequent, unbiased simulation of reentry into the additional subpocket (Replica I).

## Data availability

The coordinates and scripts generated for this study are freely available in Zenodo at the DOI: 10.5281/zenodo.10550469

## Acknowledgements

The authors gratefully acknowledge the support of the Klaus Tschira Foundation (SIMPLAIX project 3) and the provision of computing resources by the state of Baden-Württemberg through bwHPC and the German Research Foundation (DFG) through grant INST 35/1134-1 FUGG, and by the high-performance computing center, Stuttgart, Germany (HLRS; Project Dynathor). The NAMD software was developed by the Theoretical and Computational Biophysics Group in the Beckman Institute for Advanced Science and Technology at the University of Illinois at Urbana-Champaign. The authors thank Stefan Richter for assistance with technical aspects of the computations. The authors thank Sophia Ber for helpful discussions. The authors thank Nicholas Michelarakis and Florian Franz for technical advice on optimizing simulation performance.

## Author contributions

RCW conceived and supervised the study, SBH, JT and GM designed and performed the calculations. SBH, JT and RCW analysed the data and wrote the manuscript.

