## Supplementary Material for "Multiresolution molecular dynamics simulations reveal the interplay between conformational variability and functional interactions in membrane-bound cytochrome 2B4"

**Table S1**. Structural parameters characterizing the globular domain conformation and position with respect to the membrane in the CG and AA MD simulations.

| Parameters | CG representative structures (Converted to all-atom detail) | | | Final value of AA MD (average over last 50 ns) | | |
|---|---|---|---|---|---|---|
| | closed | open | Altern-ative open | closed | open | Altern-ative open |
| Angle α (°) | 91.6 | 66.0 | 67.3 | 102.2±3.3 | 62.0±2.8 | 69.7±3.5 |
| Angle β (°) | 108.3 | 75.3 | 100.4 | 131.5±4.6 | 73.7±4.1 | 97.6±3.8 |
| Heme tilt angle (°) | 46.2 | 36.7 | 41.2 | 67.9±4.0 | 39.9±4.3 | 39.5±4.8 |
| TM helix tilt angle (°) | 13.8 | 7.2 | 17.4 | 10.1±4.8 | 14.1±6.1 | 15.8±4.9 |
| D:CYP-mem (Å) | 56.4 | 50.0 | 44.5 | 40.8±2.1 | 41.7±1.3 | 43.7±1.2 |
| D:BC loop-mem (Å) | 48.4 | 27.2 | 27.5 | 36.8±1.9 | 15.8±1.3 | 26.0±1.2 |
| D:FG loop-mem (Å) | 36.3 | 32.3 | 28.6 | 22.2±1.9 | 25.3±1.8 | 28.0±1.8 |
| D:linker-mem (Å) | 39.4 | 37.8 | 24 | 22.1±1.9 | 30.4±2.0 | 26.8±1.5 |
| D:A'-F' (Å) | 13.3 | 23.7 | 24.7 | 14.8±0.3 | 16.4±0.3 | 31.6±0.8 |
| D:B'-G (Å) | 8.7 | 25.3 | 26.1 | 9.0±0.2 | 20.6±0.6 | 22.8±0.6 |
| D:B'-Cterm (Å) | 13.6 | 22.6 | 24.4 | 14.5±0.3 | 21.8±0.6 | 24.8±0.8 |

**Table S2**. Results of the CAVER analysis of the last 100 ns of the conventional MD simulations of 'closed' CYP 2B4. Snapshots were analysed at 20 ps intervals.

| | |
|---|---|
| **Cluster ID**[a] | 1 |
| **# tunnels belonging to cluster**[b] | 4999 |
| **# snapshots with >= one tunnel** | 4999 |
| **Average bottleneck radius [Å]** | 1.416 ± 0.003 |
| **Maximal bottleneck radius [Å]** | 1.42 |
| **Average length [Å]** | 22.396 ± 0.606 |
| **Average tunnel curvature**[c] | 1.378 ± 0.035 |
| **Tunnel priority calculated by averaging tunnel throughputs over all snapshot**[d] | 0.56615 |
| **Average tunnel throughput**[e] | 0.56626 ± 0.01111 |
| **Average upper error bound of bottleneck radius estimation [Å]** | 0.007 ± 0.003 |
| **Average upper error bound of tunnel profile radii estimation [Å]** | 0.045 |
| **Maximal upper errors bound of bottleneck radii estimation [Å]** | 0.027 |
| **Maximal upper error bound of tunnel radii estimation [Å]** | 0.051 |

[a]ID from cluster ranking, here only one cluster was detected.
[b]Each individual observation of a cluster in any frame corresponds to a unique tunnel.
[c]Curvature is defined as *length/distance* with distance being the shortest possible distance between the start- and end-points of the tunnel.
[d]Priority is the averaged sum of throughputs over all snapshots.
[e]Throughput is a measure of the internal, unitless cost-function, utilized by CAVER and computed from the tunnel length & -diameter defined as *throughput* = $e^{-cost}$ with the cost function defined in the work of Chovancova and others[23].

**Table S3**. Results of the AquaDuct analysis and clustering of the last 100 ns of the conventional MD simulations of the closed and open CYP 2B4.

| **System** | **Cluster[a]** | **Size [#inlets][b]** | **#Incom-ing [#inlets]** | **#Out-going [#inlets]** | **Area _D100 [$Å^2$][c]** | **Area _D95 [$Å^2$][c]** | **Area _D90 [$Å^2$][c]** | **Area _D80 [$Å^2$][c]** | **Area _D70 [$Å^2$][c]** | **Area _D60 [$Å^2$][c]** | **Area _D50 [$Å^2$][c]** |
|---|---|---|---|---|---|---|---|---|---|---|---|
| Closed | outliers | 14 | 7 | 7 | 3858.18 | 3304.63 | 3222.29 | 3027.03 | 2704.69 | 2288.23 | 1695.26 |
| Closed | 1 | 42 | 20 | 22 | 345.93 | 312.35 | 291.58 | 241.56 | 182.54 | 134.92 | 91.16 |
| Closed | 2 | 9 | 7 | 2 | 157.77 | 157.77 | 157.77 | 157.77 | 157.46 | 152.67 | 133.48 |
| Open | outliers | 29 | 15 | 14 | 10943.26 | 7735.16 | 7568.35 | 7016.11 | 5796.24 | 3324.06 | 2509.34 |
| Open | 1 | 1901 | 955 | 946 | 1278.51 | 362.91 | 302.06 | 231.41 | 187.66 | 155.38 | 122.61 |
| Open | 2 | 1730 | 869 | 861 | 1908.7 | 452.01 | 372.17 | 287.28 | 232.69 | 185.84 | 138.58 |
| Open | 3 | 147 | 64 | 83 | 198.78 | 161.91 | 139.25 | 95.03 | 62.74 | 47.02 | 35.29 |
| Open | 4 | 57 | 27 | 30 | 342.84 | 250.47 | 209.46 | 142.37 | 110.16 | 85.59 | 65.9 |
| Open | 5 | 28 | 11 | 17 | 236.27 | 153.28 | 137.66 | 111.04 | 87.72 | 72.34 | 58.96 |
| Open | 6 | 11 | 6 | 5 | 4090.81 | 2363.8 | 2258.54 | 1226.1 | 452.93 | 116.73 | 39.85 |
| Open | 7 | 8 | 6 | 2 | 101.18 | 87.98 | 83.78 | 77.6 | 70.94 | 61.23 | 50.35 |

[a]AquaDuct internal classification, no relation to the nomenclature established by Cojocaru et al.[12]
[b]Number of entry/exit points on the convex hull.
[c]Cluster area on the convex hull at different density-isovalues computed from a Kernel Density Approximation on the inlet-points. D100 corresponds to the estimated area at 100 % probability density, D50 to the area at 50 % density, etc.

**Table S4**. Results of docking BZP to crystal structures of CYP 2B4 using Autodock Vina. Poses are shown in Supplementary Figure S7.

| **Conform-ational state** | **Pose rank** | **Docking Affinity [kcal/mol]** | **Distance Fe-CH3 (Å)** | **Angle S-Fe-CH3 (°)** |
|---|---|---|---|---|
| Closed | 1 | -9.8 | 9.4 | 169.9 |
| Closed | 2* | -8.5 | 6.7 | 141.3 |
| Closed | 3 | -8.4 | 10.7 | 172.4 |
| Closed | 4 | -6.9 | 23.4 | 135.9 |
| Closed | 5 | -6.8 | 25.1 | 130.7 |
| Open | 1 | -8.2 | 10 | 124 |
| Open | 2 | -8.1 | 8.4 | 123.5 |
| Open | 3 | -7.9 | 9.9 | 127.2 |
| Open | 4 | -7.8 | 9 | 125.4 |
| Open | 5 | -7.7 | 9.2 | 122.6 |
| Open | 6 | -7.6 | 8.4 | 138.4 |
| Open | 7 | -7.5 | 7.9 | 117.9 |
| Open | 8* | -7.5 | 6.9 | 126.2 |
| Open | 9 | -7.3 | 7.8 | 143.2 |
| Open | 10 | -7.3 | 4.6 | 152.8 |

*Selected poses used in subsequent MD simulations.

**Table S5**. Ligand egress statistics from RAMD simulations

| Pathway | BZP, closed: Occurrence (%) | NZP, closed: Occurrence (%) | BZP, open: Occurrence (%) | NZP, open: Occurrence (%) | BIF, open: Occurrence (%) |
|---|---|---|---|---|---|
| **Random force magnitude (kcal/mol Å$^{-1}$)** | 12 | 12 | 6 | 6 | 8 |
| **1** | 20 | 8.9 | 0 | 0 | 0 |
| **2a** | 2.2 | 17.8 | 0 | 0 | 0 |
| **2b** | 0 | 0 | 6.7 | 0 | 22.2 |
| **2c** | 4.4 | 17.8 | 53.3 | 50 | 42.2 |
| **2ac** | 0 | 0 | 6.7 | 23.3 | 0 |
| **2d** | 0 | 0 | 0 | 0 | 2.2 |
| **2e** | 48.9 | 40 | 6.7 | 0 | 11.1 |
| **2f** | 4.4 | 2.2 | 0 | 0 | 0 |
| **4** | 0 | 0 | 0 | 0 | 2.2 |
| **Water channel** | 8.9 | 4.4 | 0 | 3.3 | 0 |
| **Solvent channel** | 4.4 | 8.9 | 0 | 0 | 0 |
| **No egress*** | 6.7 | 0 | 26.7 | 23.3 | 20 |

*Trajectories were simulated for a maximal duration of 4 ns. 'No egress' was assigned if the ligand remained within the protein for the entire trajectory.

**Table S6**. List of all ligand reentry cMD simulations carried out. The key features of the initial systems and the observed outcomes are given. The open conformation of CYP 2B4 was used for the simulations.

| Ligand | Exit pathway in preceding RAMD simulation | Reentry observed? | Reentry trajectory length (ns) | Reentry or detach-ment time (ns) | Comment on ligand movement |
|---|---|---|---|---|---|
| BIF | 2c | no | 118 | 60 | Detachs via G-helix. |
| BIF[1] | 2c | yes | 770 | ~50 | Reenters to additional subpocket in F/G cassette region. |
| BIF[2] | 2c | yes | 834 | ~50-100 | Reenters to additional subpocket in F/G cassette region. |
| BIF | 2d/f | no | 660 | - | Moves into memb-rane and associates to BC-loop. |
| BIF | 2d/f | no | 147 | 55 | Detachs into membrane via linker domain & TM-helix. |
| BIF | 2d/f | no | 677 | - | Moves into membrane and associates to BC-loop. |
| BZP | 2c | no | 228 | 78 | Rapidly detachs via C- and I-helices. |
| BZP | 2c | no | 170 | 92 | Rapid detachs via G-helix. |

[1]Replica I in Fig. 8; [2]Replica II in Fig. 8

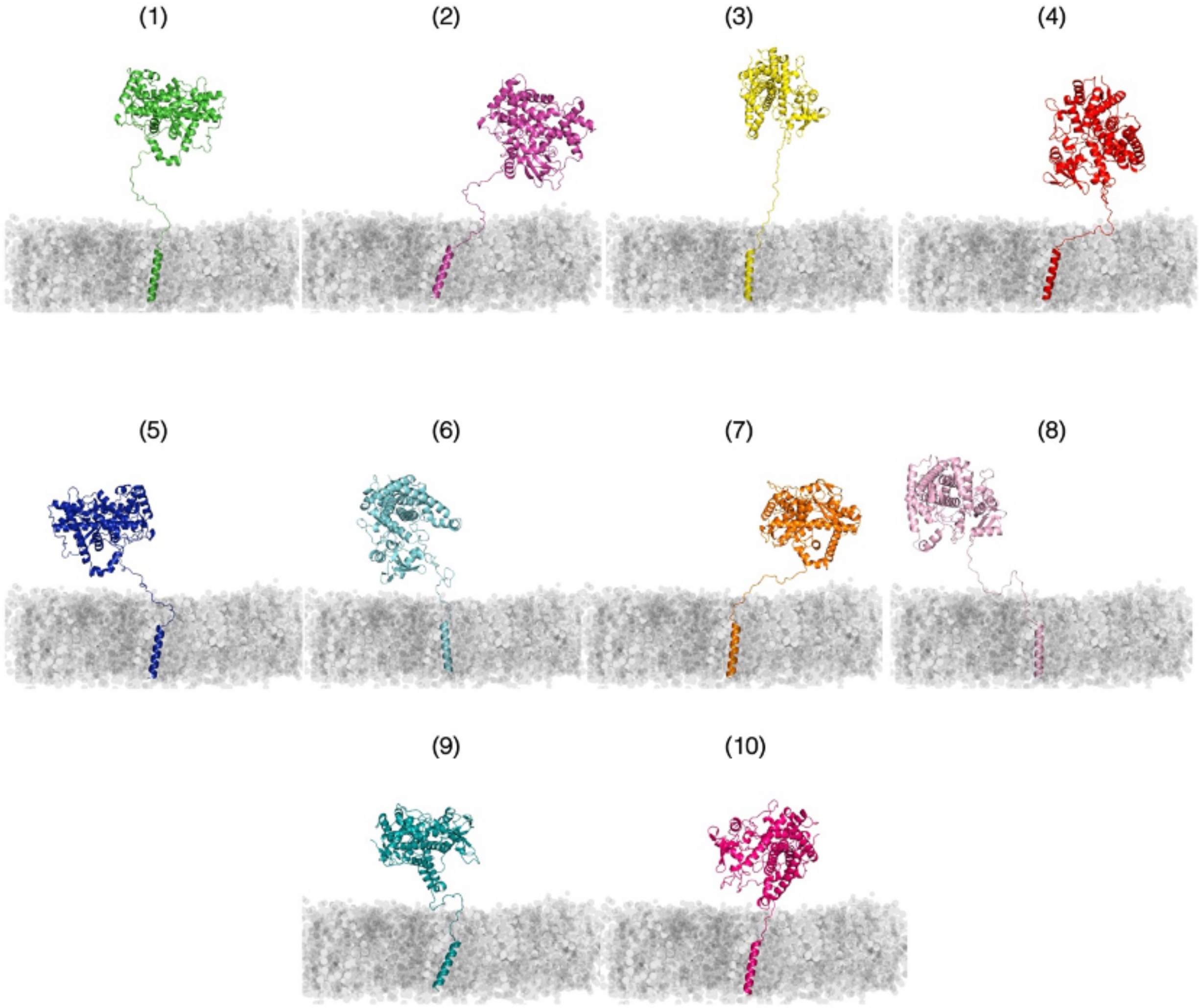


**Figure S1**. Modelled structures of CYP 2B4 with 10 different initial orientations of the globular domain in the open conformation with respect to the POPC lipid bilayer. Structures 1-10 each represent a starting structure for the 10 CG MD simulations. A similar diversity of structures was generated for the CG MD simulations with the closed conformation of CYP 2B4.

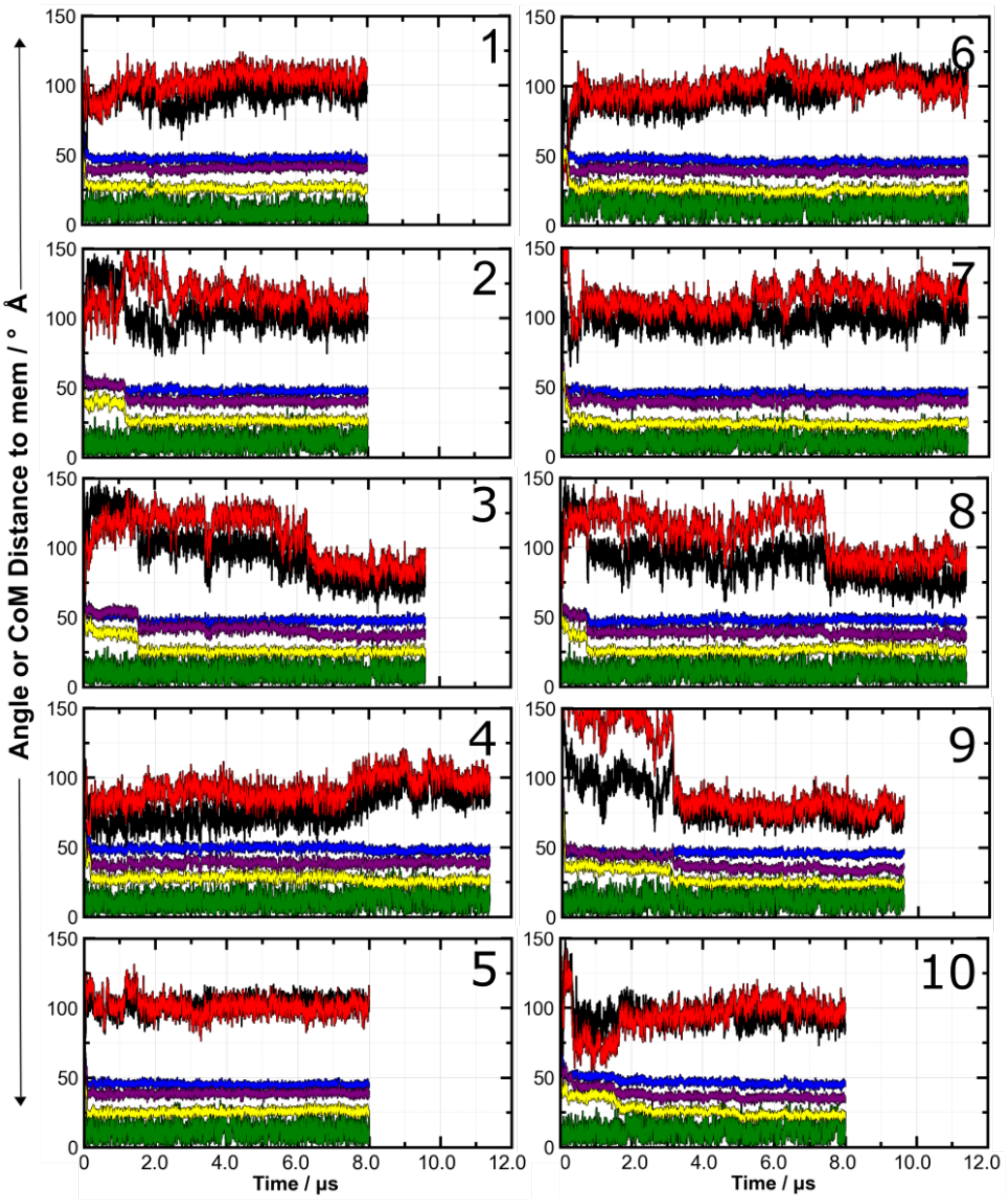


**Figure S2**. Time evolution of the positioning of the CYP 2B4 globular domain and the TM helix with respect to the lipid bilayer in 10 replica CG simulations (1-10) starting with the crystal structure of the globular domain in a closed conformation (PDB ID: 1SUO) and randomized positions of the globular domain with respect to the lipid bilayer. Color scheme: α angle (black), β angle (red), TM helix tilt angle (green), axial distances of the center of mass of the lipid membrane to the center of mass of the globular domain (blue), BC loop (purple), and FG loop (yellow).

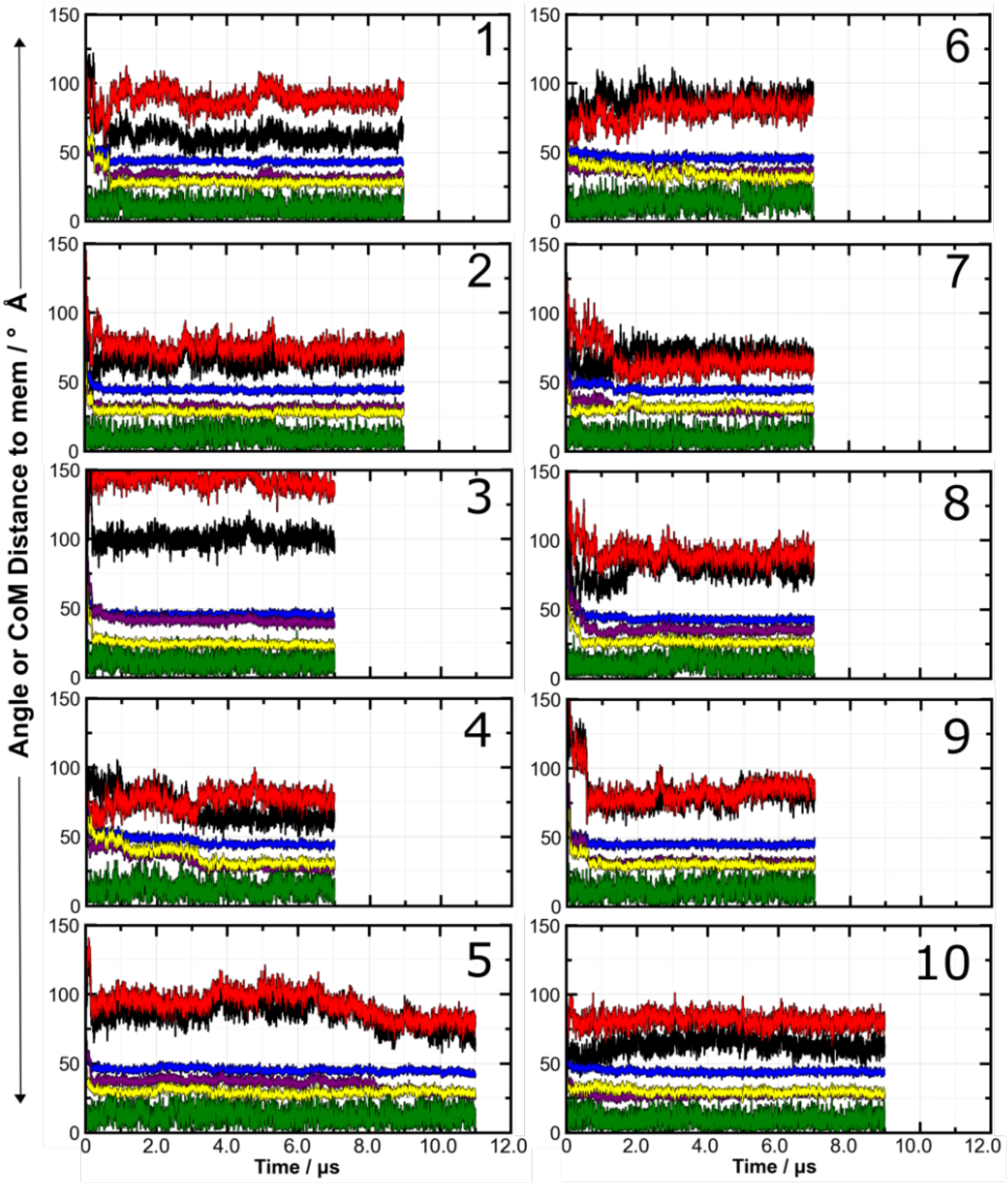


**Figure S3**. Time evolution of the positioning of the CYP 2B4 globular domain and the TM helix with respect to the lipid bilayer in 10 replica CG simulations (1-10) starting with the crystal structure of the globular domain in an open conformation (PDB ID: 1PO5) and randomized positions of the globular domain with respect to the lipid bilayer. Color scheme: α angle (black), β angle (red), TM helix tilt angle (green), axial distances of the center of mass of the lipid membrane to the center of mass of the globular domain (blue), BC loop (purple), and FG loop (yellow).

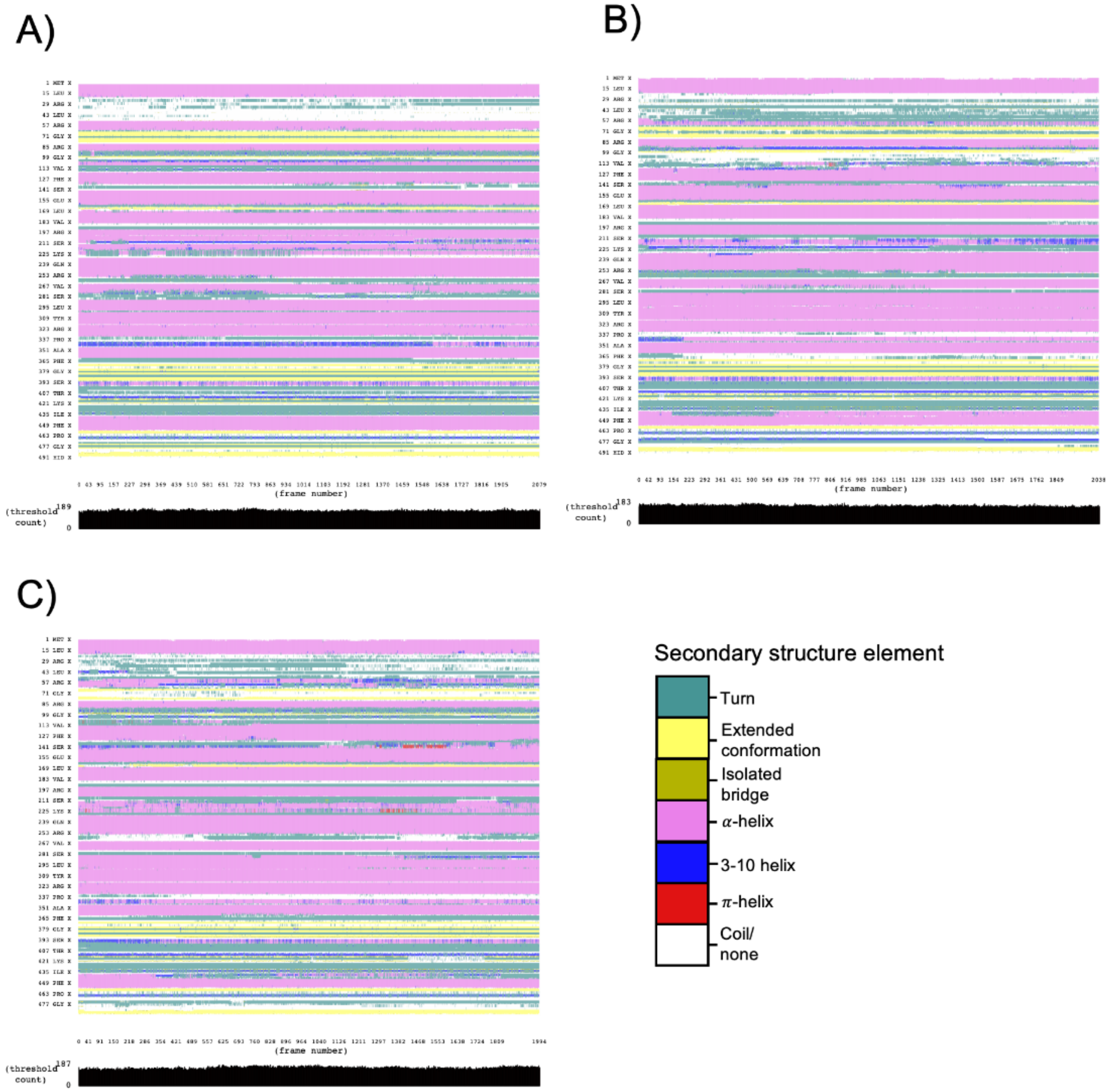


**Figure S4**. Residue-wise secondary structure classification shown along the three trajectories of CYP 2B4 in a membrane: A) closed, B) open, and C) alternative-open conformations.

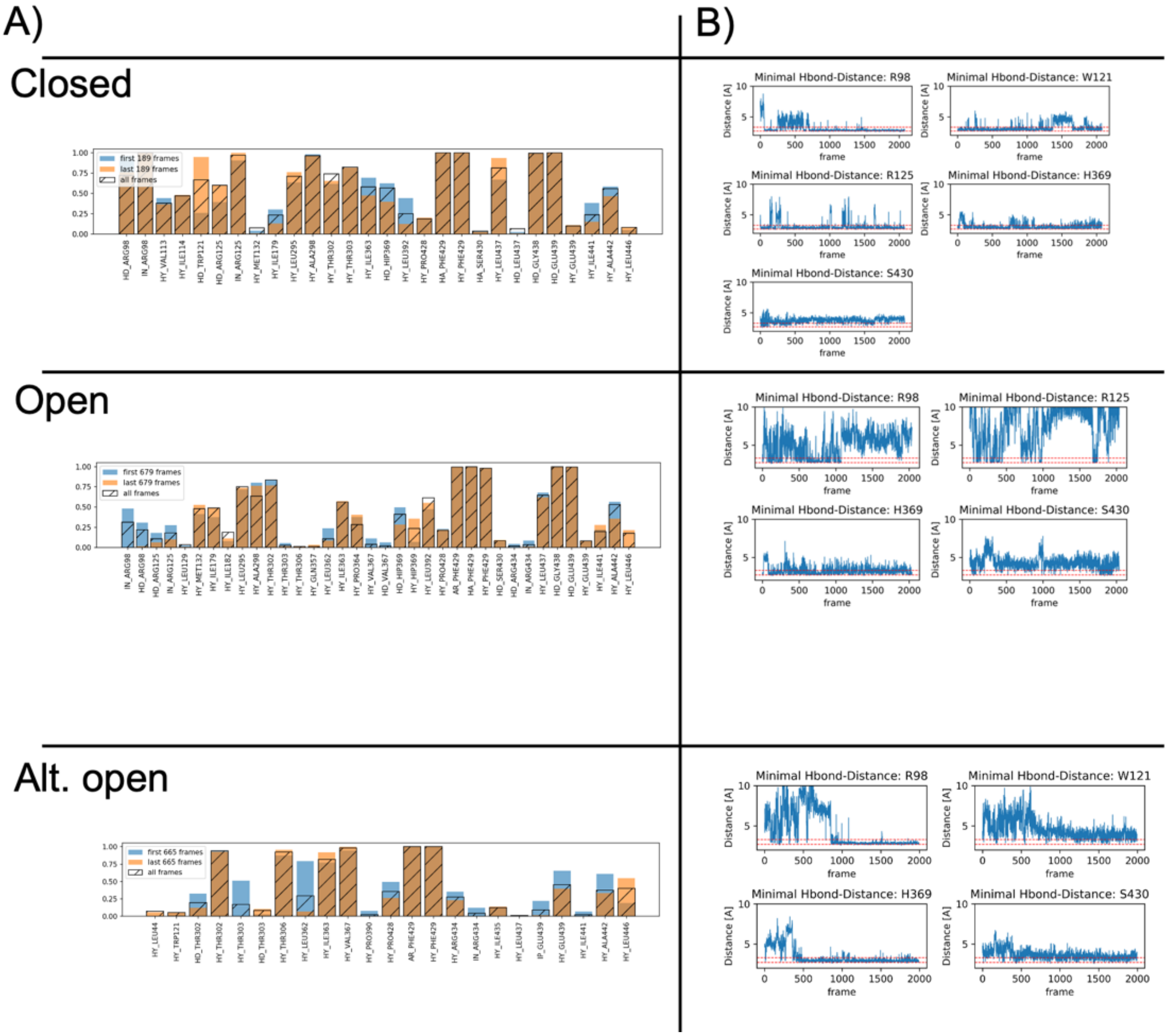


**Figure S5.** Heme-protein interaction patterns of the three CYP 2B4-membrane systems. A) MD-IFP results for the entire combined heme-cysteine residue. Blue bars indicate occupancies in the first half of the trajectory, orange bars indicate interactions in the last half. Black, transparent bars indicate occupancies over the entire trajectory. B) Hydrogen-bonding distances between the propionate oxygens and their interaction partners (interacting in more than 5 % of the analyzed frames). The red, dashed lines are at distances of 2.7 and 3.3 Å to indicate h-bond formation.

| BC-loop region | | | | DEKR | FWLVIMP | Total |
|---|---|---|---|---|---|---|
| 2B4: | 97 | GRGKIAVVDPI---FQGYGV-IFANG | 118 | 3 | 9 | 22 |
| 2C19: | 96 | GRGHFPLAERA---NRGFGI-VFSNG | 117 | 4 | 7 | 22 |
| 2C9: | 96 | GRGIFPLAERA---NRGFGI-VFSNG | 117 | 4 | 8 | 22 |
| 2E1: | 98 | GRGDLPAF-HA---HRDRGI-IFNNG | 119 | 5 | 6 | 21 |
| 2D6: | 100 | DRPPVPITQILGFGPRSQGVFLARYG | 126 | 4 | 12 | 26 |
| 1A2: | 107 | GRPDLYTSTLIT--DGQSLTFSTDSG | 131 | 4 | 6 | 24 |
| 3A4: | 104 | NRRPFGPV---G--FMKSAI-SIAED | 123 | 5 | 7 | 21 |
| 1A1: | 105 | GRPDLYTFTLIS--NGQSMSFSPDSG | 128 | 3 | 8 | 24 |
| 17A1: | 95 | GRPQMATLDIAS--NNRKGIAFADSG | 118 | 5 | 6 | 24 |

**Figure S6**. Part of a multiple sequence alignment of mammalian CYPs showing the region of the BC loop. Charged polar residues are highlighted in red and their residue count is listed under the DEKR column. The hydrophobic residue count is in the next column labeled FWLVIMP. The total number of residues in this region is given in the last column. CYP 2B4 has a more hydrophobic BC loop region than most human drug metabolizing CYPs.

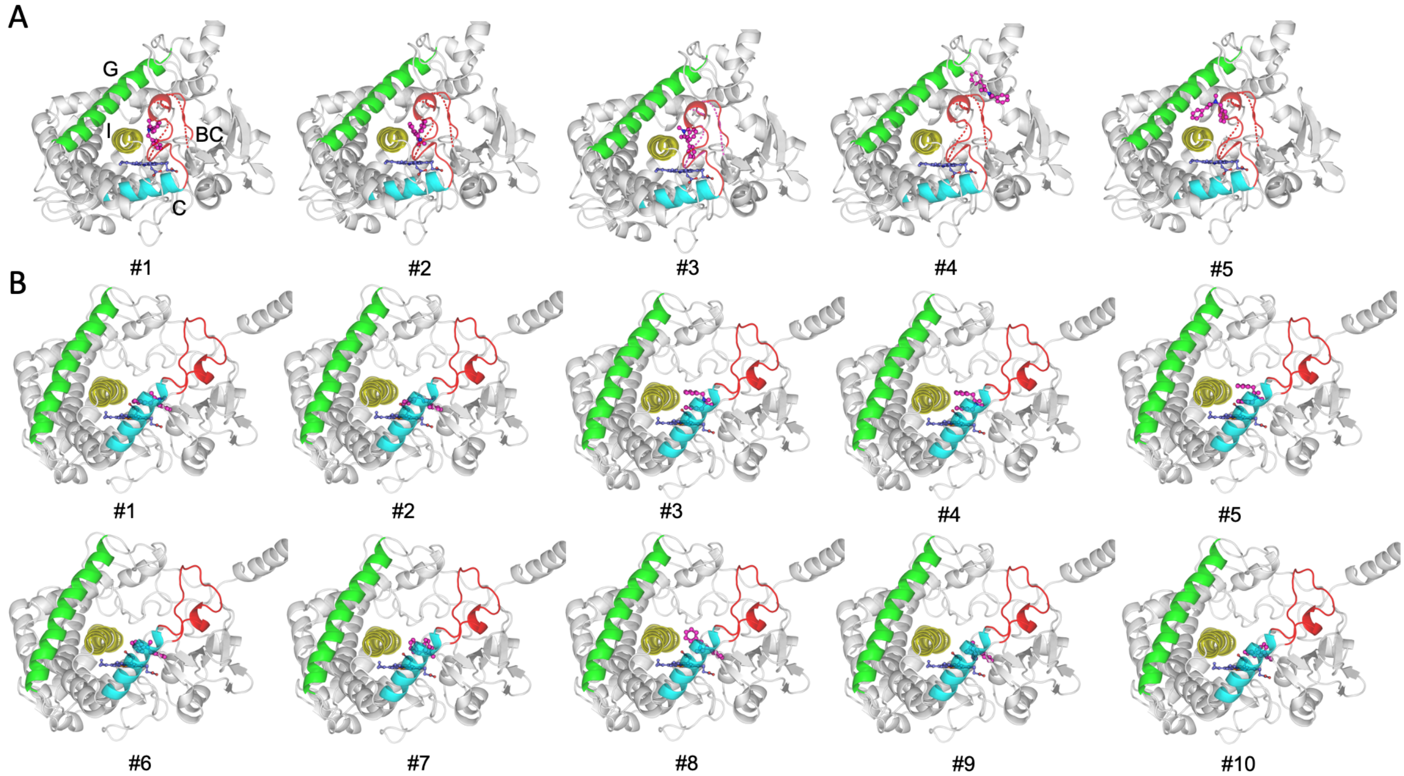


**Figure S7**. Docking poses of BZP to (A) closed and (B) open CYP 2B4. BZP is shown in magenta ball-and-stick representation. Regions with residues selected to be flexible during docking are displayed both in their original conformation as continuous ribbons and as dashed ribbons indicating the reoriented conformation. The poses are ranked according to the docking scores given in Table S2, with #1 having the best docking score.

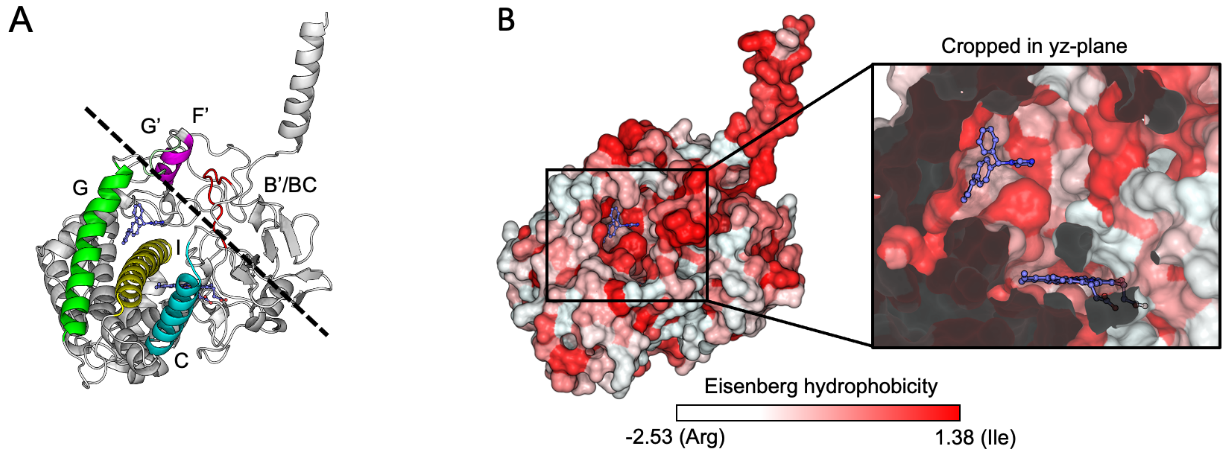


**Figure S8**. Position (A) and surface hydrophobicity (B) of the additional transient binding site for bifonazole identified in the re-entry AA MD simulations. The last frame of the Replica I AA MD simulation is shown. The tighter packing around BIF's diphenyl moiety and higher density of hydrophobic residues in the transient subpocket in comparison to the canonical binding site is visible in the inset surface representation cropped in the yz-plane.

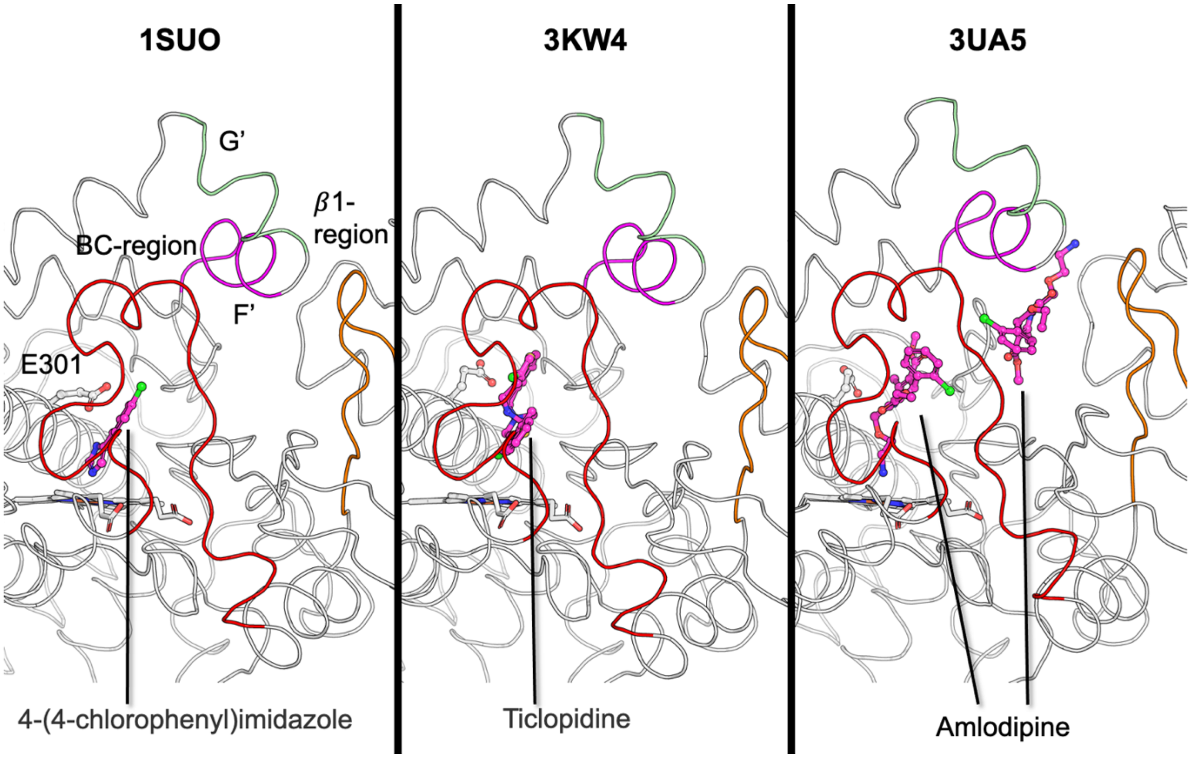


**Figure S9**. Structural differences in the conformations of CYP 2B4 upon binding of different ligands. The 1SUO and 3KW4 structures are closed (as observed for many other CYP 2B4-ligand crystal structures) but differ in the heme iron coordination of the ligand. Notably, Ticlopidine (shown with the two coordinate sets modeled in the crystal structure) extends into a hydrophobic pocket formed when the E301 sidechain rotates away from the binding site. On the other hand, in the 3UA5 structure, simultaneous binding of two Amlodipine molecules induces a subtle opening by rearrangements of the BC-loop and β1 regions and the F'- and G'-helices. The intermediately open structure induced by Bifonazole (PDB-ID: 2BDM) has much greater structural changes and is shown in Fig. 8. Similar structural changes are observed in other CYP 2B4-ligand complexes (PDB-ID 2R1B, 3G5N, 3G93). The unliganded 1PO5 structure has a wider open active site (compare Fig. 1A and Fig. 1B).
